# Cortical distribution of neurofilaments associates with pathological hallmarks and MRI measures of atrophy and diffusivity in Parkinson’s disease

**DOI:** 10.1101/2022.08.10.503440

**Authors:** Irene Frigerio, Max A Laansma, Chen-Pei Lin, Emma JM Hermans, Maud MA Bouwman, John GJM Bol, Yvon Galis-de Graaf, Dagmar H Hepp, Annemieke JM Rozemuller, Frederik Barkhof, Wilma DJ van de Berg, Laura E Jonkman

**Affiliations:** Amsterdam UMC location Vrije Universiteit Amsterdam, Department of Anatomy and Neurosciences, Section Clinical Neuroanatomy and Biobanking, De Boelelaan 1118, Amsterdam, Netherlands; Amsterdam Neuroscience, Neurodegeneration, Amsterdam, The Netherlands; Amsterdam Neuroscience, Brain imaging, Amsterdam, The Netherlands; Amsterdam UMC location Vrije Universiteit Amsterdam, Department of Neurology, De Boelelaan 1117, Amsterdam, Netherlands; Amsterdam UMC location Vrije Universiteit Amsterdam, Department of Pathology, De Boelelaan 1117, Amsterdam, Netherlands; Amsterdam UMC location Vrije Universiteit Amsterdam, Department of Radiology and Nuclear Medicine, De Boelelaan 1117, Amsterdam, Netherlands; University College London, Institutes of Neurology and Healthcare Engineering, London, United Kingdom

**Author notes:** Correspondence to: Irene Frigerio, De Boelelaan 1118, 1081 HV, Amsterdam, the Netherlands.

**Keywords:** neurofilament, NfL, axonal degeneration, Parkinson’s disease, Parkinson’s disease dementia, dementia with Lewy Bodies, cortical thickness, cortical atrophy, mean diffusivity

## Abstract

Increased neurofilament levels in biofluids, commonly used as a proxy for axonal degeneration in the brain, have been found in Parkinson’s disease (PD), PD with dementia (PDD) and dementia with Lewy bodies (DLB), and correlate with MRI biomarkers. The aim of the current study was to unravel the regional distribution of cortical neurofilaments and their association with pathology load and MRI measures of atrophy and diffusivity in the post-mortem brain.

Using a within-subject post-mortem MRI-pathology approach, we included 9 PD, 12 PDD/DLB and 18 age-matched control donors. Cortical thickness and mean diffusivity (MD) metrics were extracted respectively from 3DT1 and DTI at 3T *in-situ* MRI. After autopsy, pSer129 alpha-synuclein (pSer129-αSyn), p-tau, and amyloid-beta, together with neurofilament light-chain (NfL) and phosphorylated neurofilament medium- and heavy-chain (p-NfM/H) immunoreactivity were quantified in 7 cortical regions, and studied in detail with confocal-laser scanning microscopy. The correlations between MRI and pathological measures were studied using linear mixed models.

Compared to controls, p-NfM/H immunoreactivity was increased in all cortical regions in PD and PDD/DLB, whereas NfL immunoreactivity was mainly increased in the parahippocampal and entorhinal cortex in PDD/DLB. NfL-positive neurons showed degenerative morphological features and axonal fragmentation. Increased p-NfM/H correlated with p-tau load, and NfL correlated with pSer129-αSyn but more strongly with p-tau load in PDD/DLB. Lastly, neurofilament immunoreactivity correlated with cortical thinning in PD and with increased cortical MD in PDD/DLB.

Taken together, increased neurofilament immunoreactivity suggests underlying axonal injury and neurofilament accumulation in morphologically altered neurons with increasing pathological burden. Importantly, we demonstrate that such neurofilament markers at least partly explain MRI measures that are associated with the neurodegenerative process.

## Introduction

Parkinson’s disease (PD) is a common neurodegenerative disease with a heterogeneous clinical presentation, which is diagnosed when bradykinesia and tremor and/or rigidity are present [56]. Cognitive impairment is common in PD, leading to Parkinson’s disease dementia (PDD) in up to 80% of PD patients [1, 22, 23, 37]. However, when patients develop dementia before or within one year of motor symptom onset, the disease is diagnosed as dementia with Lewy bodies (DLB) [49]. Pathologically, PD, PDD and DLB are defined as synucleinopathies [38], since they are characterized by the accumulation of alpha synuclein (αSyn) in the form of Lewy bodies (LBs) and Lewy neurites (LNs), which are abundant in the cortex of demented patients [36]. In addition, PDD and DLB patients frequently show Alzheimer’s disease (AD) co-pathology, such as amyloid-beta (Aβ) plaques and phosphorylated-tau (p-tau) neurofibrillary tangles (NFT) [31, 47]. The pathological hallmarks of PDD and DLB largely overlap, and only few neuropathological differences between the two have been described [29, 49, 64].

In addition to pathology load, there is growing evidence for the presence of axonal degeneration in PD, PDD and DLB [7, 41]. Neurofilaments in cerebrospinal fluid (CSF) and blood are commonly used as a proxy of axonal degeneration in many neurodegenerative disorders [7, 41]. Neurofilaments are neuronal-specific proteins which consists of neurofilament light (NfL), medium (NfM) and heavy chain (NfH) [41]. CSF and blood NfL levels have been shown to be elevated in PD, and more abundantly in PDD and DLB [44, 45, 50, 54, 67], and to associate with higher levels of CSF αSyn and p-tau biomarkers [28] as well as with measures of cognitive decline [7, 9, 44, 45, 53, 67]. However, the distribution pattern of NfL and phosphorylated neurofilament medium and heavy chain (p-NfM/H) in cortical brain regions, and to which extent these neurofilaments are related to pathological accumulation in PD, PDD and DLB is yet to be determined.

Atrophy and altered microstructural integrity of the cortex can be captured with MRI outcome measures such as cortical thickness and mean diffusivity (MD), and may be a proxy of underlying neurodegeneration [57]. While PDD and DLB patients show more pronounced cortical atrophy and increased cortical MD compared to unimpaired elderly [8, 12, 16, 65], PD patients show only subtle changes [42, 58, 61]. Cortical atrophy and increased cortical MD were shown to correlate with plasma NfL in several cortical regions in *de-novo* PD [59]. However, little is known about the relation between regional distribution of neurofilaments and MRI-derived microstructural measures in the brain in PD, PDD and DLB.

The aim of the current study was to unravel the regional distribution of cortical neurofilaments, and their association with pathology load and MRI measures of cortical thickness and diffusivity in the brain of clinically-defined and pathologically-confirmed PD, PDD/DLB and control brain donors, using a within-subject post-mortem MRI-pathology approach [40]. Results of this study will increase knowledge on the regional distribution of neurofilaments in the cortex in PD, PDD and DLB and to what extent this is related to pathological accumulation, and reflected by cortical thickness and diffusivity measures, thereby contributing to the understanding of the pathological underpinnings of MRI.

## Materials and methods

### Donor inclusion

In collaboration with the Netherlands Brain Bank (NBB; http://brainbank.nl) we included 21 Lewy Body disease donors who could be further subdivided into 9 PD, 7 PDD and 5 DLB based on clinical presentation [23, 49, 56]. Age at diagnosis (at symptoms onset) and disease duration (age at death minus age at diagnosis) were extracted from the clinical files of PD(D)/DLB donors. Neuropathological diagnosis was confirmed by an expert neuropathologist (A.R.) and performed according to the international guidelines of the Brain Net Europe II (BNE) consortium (https://www.brainbank.nl/about-us/brain-net-europe/) [3–5]. Additionally, 18 age- and sex-matched and pathologically confirmed controls with no records of neurological symptoms at diagnosis, were selected from the Normal Aging Brain Collection Amsterdam (NABCA; http://nabca.eu) [40]. All donors signed an informed consent for brain donation and the use of material and clinical information for research purposes. The procedures for brain tissue collection of NBB and NABCA have been approved by the Medical Ethical Committee of Amsterdam UMC, Vrije Universiteit Amsterdam. For donor characteristics, see Supplementary Table 1, online resource 1.

### Post-mortem *in-situ* MRI acquisition

Post-mortem *in-situ* (brain in cranium) MRI scans were acquired according to a previously described pipeline [40] (**Fig. 1**). Briefly, scans were acquired on a 3T scanner (Signa-MR750, General Electric Medical Systems, United States) with an eight-channel phased-array head-coil. The following pulse-sequences were performed for all subjects: i) sagittal 3D T1-weighted fast spoiled gradient echo (repetition time [TR] = 7 ms, echo time [TE] = 3 ms, flip angle = 15°, 1-mm-thick axial slices, in-plane resolution = 1.0 x 1.0 mm^2^); ii) sagittal 3D fluid attenuation inversion recovery (FLAIR; TR = 8000 ms, TE = 130 ms, inversion time [TI] = 2000-2500 ms, 1.2-mm-thick axial slices, in-plane resolution = 1.11 x 1.11 mm^2^), with TI corrected for post-mortem delay; iii) diffusion-weighted imaging (DWI) axial 2D echo-planar imaging with diffusion gradients applied in 30 non-collinear directions, TR/TE= 7400/92 ms, slices thickness of 2.0 mm, in-plane resolution = 2.0 x 2.0 mm^2^, and b = 1000 s/mm^2^. To allow for offline distortion correction of the images, 5 b0 images were acquired using the same sequence parameters.

**Fig. 1.**
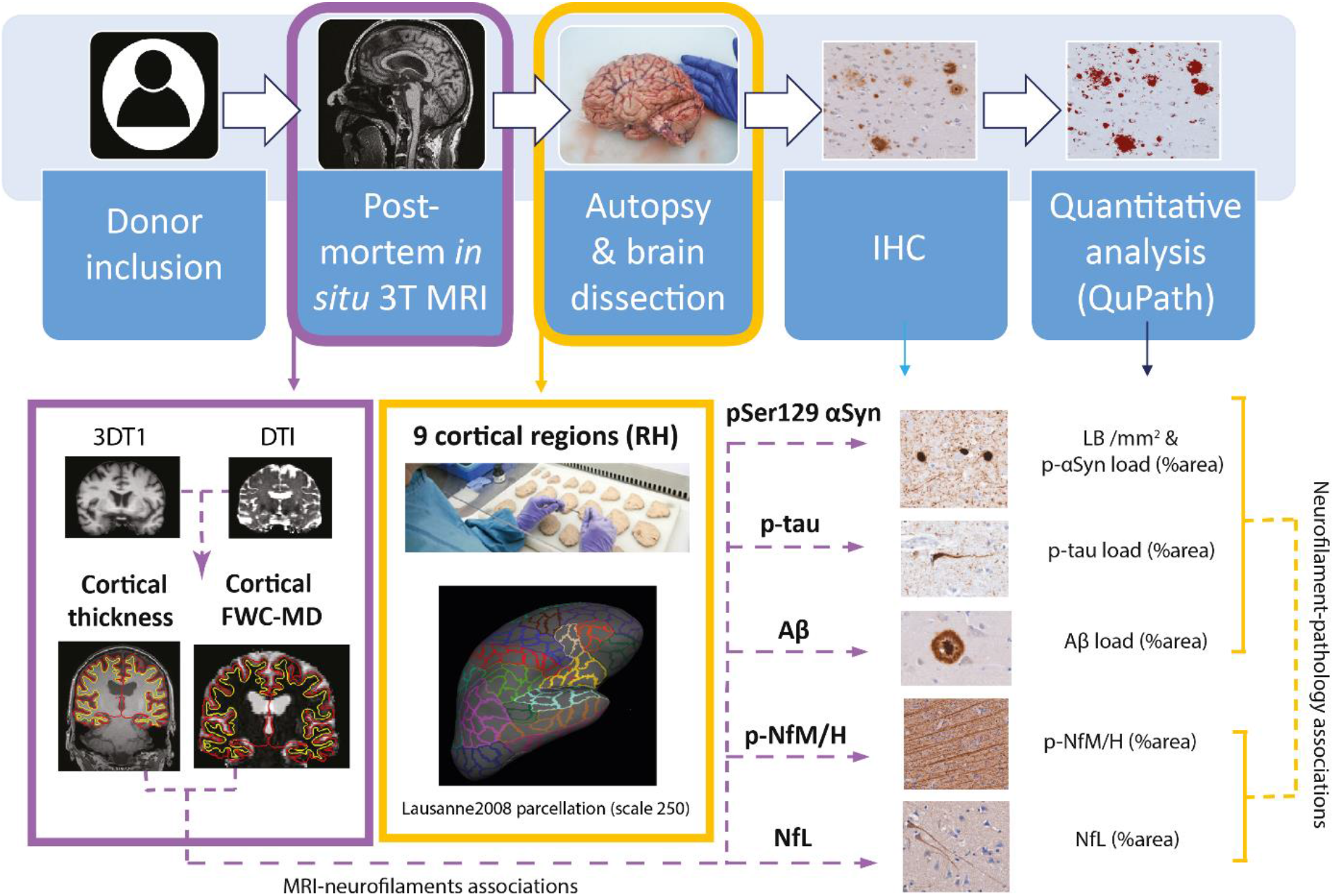
Workflow. After donor inclusion, post-mortem *in-situ* 3DT1 and DTI were collected [40], from which cortical thickness was derived with Freesurfer [21] and free-water corrected cortical mean diffusivity (FWC-MD) was derived with FSL [39], DIPY [25, 33] and Freesurfer [20] (purple box). After the MRI scan, autopsy was performed, and 9 cortical regions were selected from the right hemisphere (RH) for further analysis. To match these regions to the MR images, regions of interest were selected from the Lausanne atlas [20, 27] (yellow box). Brain tissue was processed for immunohistochemistry against phosphorylated Ser129 α-synuclein (pSer129-αSyn, clone EP1536Y), phosphorylated-tau (p-tau, clone AT8), amyloid β (Aβ, clone 4G8), neurofilament light chain (NfL, amino acid sequence 1-376) and phosphorylated neurofilaments medium and heavy chains (p-NfM/H, clone SMI312), which were quantified using QuPath [10]. The correlations between neurofilament immunoreactivity and pathology load, and between neurofilament immunoreactivity and MRI outcome measures were investigated via linear mixed models (yellow and purple dashed arrows, respectively). Legend: *Aβ: amyloid beta; FWC-MD: free water corrected mean diffusivity; IHC: immunohistochemistry; LB: Lewy Body; NfL: neurofilament light chain; pSer129-αSyn: phosphorylated Ser129 alpha synuclein; p-NfM/H: phosphorylated neurofilament medium and heavy chain; p-tau: phosphorylated tau; RH: right hemisphere*

### MRI analysis

#### Cortical thickness and brain volume assessment

To minimize the impact of age-related white matter abnormalities (e.g. vascular change) on automated segmentations, the 3D-T1 images were lesion-filled [60] as previously described [24, 46]. Image processing was performed using Freesurfer, version 6.0 (http://surfer.nmr.mgh.harvard.edu) [21]. For each subject, 9 cortical regions of interest (ROIs) in the right hemisphere were selected from the Lausanne atlas parcellation [20, 27], to closely match the location of the cortical regions dissected at autopsy. Details on the Lausanne atlas labels corresponding to the ROIs in the present study can be found in Supplementary Table 2 (online resource 1), and included the superior frontal gyrus, anterior and posterior cingulate gyrus, anterior insula, middle temporal gyrus, superior parietal gyrus, entorhinal cortex, parahippocampal and fusiform gyrus. Cortical thickness was measured as the perpendicular distance from the grey/white matter boundary to the corresponding pial surface. Quality of parcellations was assessed using the ENIGMA guidelines for quality control (see http://enigma.ini.usc.edu/protocols/imaging-protocols/). Furthermore, post-mortem normalized brain volume, and normalized grey and white matter volumes were obtained from the T1-weighted images using Structural Image Evaluation, using Normalisation, of Atrophy (SIENAX) (part of FSL 5.0.9; http://fsl.fmrib.ox.ac.uk/), which estimates brain tissue volume normalized for skull size [39].

#### DTI pre-processing and free water correction

Diffusion tensor imaging (DTI) and reference scans of opposite phase encodings (acquired along anterior-posterior and posterior-anterior directions) were collected for 9 out of 9 PD donors, 11 out of 12 PDD/DLB donors, and 14 out of 18 control donors (see Supplementary Table 1, online resource 1). DTI was first corrected for susceptibility distribution and eddy current induced geometric distortion using FSL (Eddy and topup) [39] and fitted for single tensors diffusion maps, deriving MD [19], which is a good (bio)marker for grey matter microstructure, where diffusion does not conform to a specific direction [18]. To avoid underestimation by partial volume effects of the CSF, a bi-tensor model for free water correction was performed using an open resource, python-based script of DIPY, which was shown to be plausible for using single shell DWI acquisitions to derive free-water corrected MD (FWC-MD) maps [25, 33]. Using Freesurfer version 7.1.1 [21], we performed a within-subject registration of T1-weighted to FWC-MD images. With this registration, the cortical thickness ROIs were registered to the FWC-MD map and corrected for overestimation into the white matter, by applying a threshold at 50% probability and a limit to the cortical ribbon mask, after which regional cortical FWC-MD values obtained. Quality of parcellations was manually assessed (I.F.) and, when necessary, poorly parcellated regions were manually corrected or excluded from statistical analysis (i.e. entorhinal cortex of 1 PD case and 3 PDD/DLB cases was excluded). For clarity, FWC-MD is referred as MD in the manuscript.

### Tissue sampling

Subsequent to MRI acquisition, brain tissue was collected at autopsy, resulting in a total post-mortem delay (interval between death and autopsy) of maximum 13 hours for all brain donors. Formalin-fixed paraffin-embedded tissue blocks (4%, four weeks fixation) from the following seven regions within the right hemisphere were obtained and processed for immunohistochemistry: superior frontal gyrus, anterior and posterior cingulate gyrus, anterior (dysgranular) insula, middle temporal gyrus, superior parietal gyrus, and hippocampus (including the entorhinal cortex, parahippocampal and fusiform gyrus, as described before) [2, 24].

### Immunohistochemistry (IHC) for quantification

Six μm thick sections from tissue blocks of the above-mentioned regions were cut and mounted on superfrost+ glass slides (Thermo Scientific, USA). All sections were stained for pSer129-αSyn (clone EP1536Y), Aβ (clone 4G8) and p-tau (clone AT8). Additionally, sections from 7 out of the 9 cortical brain regions (showing significantly higher LB count across groups, i.e. all regions except the posterior cingulate cortex and the superior parietal gyrus) were stained for neurofilament light chain (NfL, amino acid sequence 1-376) and phosphorylated neurofilament medium and heavy chains (p-NfM-H; clone SMI312) (for information on primary antibodies see Supplementary Table 3, online resource 1). Briefly, the sections were deparaffinised, immersed in buffer, and heated to 100°C in a steam cooker for 30 minutes for antigen retrieval. The sections were blocked for endogenous peroxidase using 1% hydrogen peroxide and in tris buffered saline (TBS; pH 7.4), and consequently in 3% normal donkey serum in TBS (Triton 0,5%). Primary antibodies were diluted in 1% normal donkey serum in TBS (Triton 0,1%) and incubated overnight at 4°C. Primary antibodies were detected using EnVision (Dako, Glostrup, Denmark), and visualized using 3.3’-Diaminobenzidine (DAB, Dako) with Imidazole (50 mg DAB, 350 mg Imidazole and 30 uL of H2O2 per 100 mL of Tris-HCl 30 mM, pH 7.6). In between steps, TBS was used to wash the sections. After counterstaining with haematoxylin, the sections were dehydrated and mounted with Entellan (Merck, Darmstadt, Germany).

### Pathological analysis

#### Pathology image processing

Using a whole-slide scanner (Vectra Polaris, 20x objective) images of immunostained sections were taken and quantified using QuPath 0.2.3 (https://qupath.readthedocs.io/en/0.2/) [10]. ROIs containing all cortical layers were delineated in straight areas of the cortex to avoid over- or underestimation of pathology in sulci and gyri, respectively [6]. Hippocampal sections were segmented according to the method described by Adler et al. [2], where the entorhinal cortex, parahippocampal gyrus and fusiform gyrus were delineated to match the MRI derived ROIs. Briefly, the entorhinal cortex was delineated from the end of the parasubiculum until layer IV started being visible [35]; the parahippocampal gyrus started from this point and ended at the collateral sulcus; the fusiform gyrus started at this point and ended at the inferior temporal sulcus (Supplementary Fig. 1, online resource 2). For the pSer129-αSyn, NfL and p-NfM/H stainings, the cortex was further segmented into superficial (layers I-III) and deep (layers IV-VI) cortical layers based on the haematoxylin counterstaining. For Aβ and p-tau, the same was done in the entorhinal cortex, parahippocampal and fusiform gyrus, as these are among the first regions to be affected by Alzheimer’s disease co-pathology [14, 63]. Note that the entorhinal cortex was subdivided into layers I-III (superficial) and lamina dissecans plus layers V-VI (deep) [35]. DAB immunoreactivity was quantified with *in-house* QuPath scripts, using pixel and object classifiers. The outcome measures for pSer129-αSyn were both LB count per mm^2^ (LB/mm^2^) and pSer129-αSyn %area load excluding LBs (for additional information, see Supplementary Material, online resource 3), while outcome measures for Aβ and p-tau were %area load (Supplementary Fig. 2, online resource 2). The outcome measures for neurofilaments were %area load, expressed in the text as %immunoreactivity (Supplementary Fig. 3, online resource 2).

#### Multi-label immunofluorescence and confocal microscopy for morphological characterisation of neurofilaments

To study the morphological features of neurofilaments, we performed a multi-label immunofluorescence staining in combination with 3D confocal laser scanning microscopy (CSLM) of p-NfM/H and NfL on a representative case of each group. This was done on the parahippocampal gyrus, as this was the region showing the largest neurofilament alterations between groups. Briefly, sections were deparaffinised, immersed in Tris EDTA pH 9.0, and heated to 100°C in a steam cooker for 30 minutes for antigen retrieval. The sections were blocked for endogenous peroxidase using 1% hydrogen peroxide in TBS, and consequently in 3% normal donkey serum in TBS (Triton 0.5%; pH 7.4). Primary antibodies were diluted in 1% normal donkey serum in TBS (Triton 0.1%; pH 7.4) and incubated overnight at 4°C. Primary antibodies were detected and visualized with donkey-anti-mouse Alexa 488 (ThermoFisher, Pittsburgh, USA) and donkey-anti-rabbit Alexa 594 (ThermoFisher, Pittsburgh, USA) targeting p-NfM/H and NfL, respectively. For all stainings, TBS was used to wash the sections between steps. After counterstaining with DAPI (Sigma-Aldrich, Missouri, United States), the sections were mounted with Mowiol (Sigma-Aldrich, Missouri, United States) plus anti-fading agent DABCO. Confocal imaging was performed with a Leica TCS SP8 (Leica Microsystems, Germany) using a HC PL PAO CS2 100x oil objective lens, NA 1.40 and a zoom factor of 2.0. Sections were sequentially scanned for each fluorochrome with a pulsed white light laser at different wavelengths (excitation wavelengths: DAPI at 405 nm; Alexa 488 at 499 nm; Alexa 594 at 598 nm). All signals were detected using gated hybrid detectors in counting mode. Z-stacks (Z= 6 μm; 1024×1024 pixels) were taken in the parahippocampal gyrus of a representative control, PD and DLB. After scanning, the images were deconvoluted using CMLE algorithms in Huygens Professional (Scientific Volume imaging; Huygens, The Netherlands; https://svi.nl/Huygens-Professional), and their maximum projections (ImageJ Fiji, National Institute of Health USA; https://imagej.nih.gov/ij/) or 3D surface rendering reconstructions (Imaris, Oxford instruments 2022: https://imaris.oxinst.com/) were used to represent graphically the structures of interests and their morphologies. In some cases, signal brightness was increased in ImageJ for clarity. Figures were created using Adobe Illustrator (CS6, Adobe Systems incorporated).

### Statistics

Statistical analyses were performed in SPSS 26.0 (Chicago, IL). Normality was tested, and demographics between PD, PDD/DLB and control groups were compared using parametric or non-parametric tests for continuous data, and Fisher exact test for categorical data. Linear mixed models were used for all analyses (both group differences and associations) with age (at death) and sex as covariates. LMM on cortical thickness and MD (dependent variables) were performed with post-mortem delay as covariate in additional to age and sex, as post-mortem delay might influence post-mortem MRI-derived biomarkers [13]. For LMM associations, cortical thickness metrics were transformed into z-scores for each brain region as different cortical regions inherently have different thicknesses, which may drive the association. Multiple comparison between groups were corrected with post-hoc Bonferroni, and group comparisons were expressed as percentage differences, as in the formula: %difference = [(absolute difference)/mean control]*100. Statistics at the brain region level were corrected for multiple comparisons using the false discovery rate approach (FDR), and FDR corrected p-value are expressed as p-FDR [11]. Correlation coefficients (r) of LMM associations were calculated as in the formula: r = (estimate fixed effect * standard deviation fixed effect) / standard deviation dependent variable.

## Data availability

The data that support the findings of this study are available from the corresponding author, upon reasonable request.

## Results

### Cohort description

Demographic, clinical, radiological, and pathological data of PD, PDD/DLB and non-neurological control donors are summarized in **Table 1** (and per donor in Supplementary Table 1, online resource 1). Sex, age at diagnosis, age at death, and post-mortem delay did not differ between groups, whereas disease duration was shorter in PDD/DLB compared to PD donors (*p=0.006*), due to the inclusion of DLB donors with shorter disease duration (n=5, mean ± SD, 5 ± 1.6 years). On MRI, normalized brain volume (*p=0.011*), and normalized grey matter volume (*p=0.019*), but not normalized white matter volume (*p=0.095*) were lower in PDD/DLB cases compared to controls. In terms of pathology load, the PDD/DLB group showed more abundant LB count, p-tau and Aβ load compared to controls (*LB/mm^2^: p<0.001; p-tau: p=0.015; Aβ: p=0.018*) and PD donors (*LB/mm^2^: p=0.033; p-tau: p=0.035; Aβ: p=0.033*). Moreover, LBs were more abundant in deep than superficial cortical layers (PD: *p=0.004; PDD/DLB:p=0.035*), while p-tau and Aβ were more abundant in superficial than deep cortical layers in PDD/DLB (*p-tau: p=0.046; Aβ: p<0.001*). P-tau load strongly correlated with both LB count (*r = 0.68, p<0.001*) and pSer129-αSyn load (*r = 0.60, p<0.001*) in the PDD/DLB group. For more details on regional and layer-specific distribution of pSer129-αSyn, Aβ, and p-tau load, see Supplementary Fig. 4, online resource 2. For more details on correlations between pathological hallmarks, see Supplementary Table 4, online resource 1.

**Table 1.** Donor’s characteristics. Data are mean ± standard deviation (SD). Legend: CAA: cerebral amyloid angiopathy; DLB: dementia with Lewy bodies; F: females; LB: Lewy body; M: males; NBV: normalized brain volume; NFT: neurofibrillary tangles; NGMV: normalized grey matter volume; NWMV: normalized white matter volume; PD: Parkinson’s disease; PDD: Parkinson’s disease dementia; SD: standard deviation. \**p<0.05*, \*\**p*<0.01, \*\*\**p*<0.001 when compared to controls, and ^Ϯ Ϯ^ *p*<0.01 when compared to the PD group.

|  | Control | PD | PDD/DLB |
| --- | --- | --- | --- |
| N | 18 | 9 | 12<br>(7 PDD, 5 DLB) |
| Sex M/F (%M) | 8/10 (44%) | 6/3 (67%) | 7/5 (58%) |
| Age at diagnosis<br>years, mean ± SD | - | 62 ± 7 | 68 ± 12 |
| Disease duration<br>years, mean ± SD | - | 18 ± 5 | 11 ± 6 <sup>††</sup> |
| Age at death<br>years, mean ± SD | 73 ± 8 | 80 ± 9 | 78 ± 9 |
| Post-mortem delay | 524 ± 110 | 469 ± 153 | 443 ± 107 |
| minutes, mean ± SD | | | |
| **Radiologic characteristics** |  |  |  |
| **NBV (L)** mean ± SD | 1.46 ± 0.07 | 1.43 ± 0.08 | 1.38 ± 0.07* |
| **NGMV (L)** mean ± SD | 0.74 ± 0.05 | 0.72 ± 0.04 | 0.70 ± 0.04* |
| **NWMV (L)** mean ± SD | 0.72 ± 0.03 | 0.72 ± 0.05 | 0.68 ± 0.04 |
| **Pathological characteristics** |  |  |  |
|  | **Control** | **PD** | **PDD/DLB** |
| **Braak LB stage [15] N** | 18 | 9*** | 12*** |
| 0/1/2/3/4/5/6 | 15/2/1/0/0/0/0 | 0/0/0/0/1/1/7 | 0/0/0/0/0/0/12 |
| **ABC score [51] N** | 18 | 9 | 12 |
| A 0/1/2/3 | 4/13/1/0 | 1/6/2/0 | 0/3/5/4** |
| B 0/1/2/3 | 3/15/0/0 | 0/7/2/0 | 0/5/7/0** |
| C 0/1/2/3 | 18/0/0/0 | 7/2/0/0 | 5/1/6/0** |
| **Thal phase [63] N** | 18 | 9 | 12** |
| 0/1/2/3/4/5 | 4/8/5/1/0/0 | 1/3/3/2/0/0 | 0/2/1/5/3/1 |
| *Thal et al. 2002* |  |  |  |
| **Braak NFT stage [14] N** | 18 | 9 | 12*** |
| 0/1/2/3/4/5/6 | 3/11/4/0/0/0/0 | 0/2/5/1/1/0/0 | 0/0/5/2/5/0/0 |
| *Braak et al. 2006* |  |  |  |
| **CAA type [62] N** | 18 | 9 | 12 |
| No CAA/type 1/ type 2 | 14/1/3 | 9/0/0 | 5/4/3 |

Data are mean ± standard deviation (SD). Legend: CAA: cerebral amyloid angiopathy; DLB: dementia with Lewy bodies; F: females; LB: Lewy body; M: males; NBV: normalized brain volume; NFT: neurofibrillary tangles; NGMV: normalized grey matter volume; NWMV: normalized white matter volume; PD: Parkinson's disease; PDD: Parkinson's disease dementia; SD: standard deviation. \* $p < 0.05$ , \*\* $p < 0.01$ , \*\*\* $p < 0.001$ when compared to controls, and $^{\dagger\dagger} p < 0.01$ when compared to the PD group.

### Increase in neurofilament immunoreactivity in the cortex of PD and PDD/DLB

P-NfM/H and NfL immunoreactivity were investigated in seven cortical areas. P-NfM/H staining was observed in axons, while NfL staining was seen in the neuronal soma and its processes (**Fig. 2a** and **b**). Both p-NfM/H and NfL immunoreactivity positively correlated with age (p-NfM/H: *r = 0.33, R^2^=11%, p=0.011;* NfL in superficial cortical layers: *r = 0.23, R^2^=5%, p=0.020*), and p-NfM/H was significantly higher in males compared to females (*p=0.007*) (Supplementary Fig. 5, online resource 2). P-NfM/H immunoreactivity was higher in deep compared to superficial cortical layers in all groups (controls: +38% compared to superficial layers, *p<0.001*; PD: +22%, *p<0.001*; PDD/DLB: +23%, *p<0.001*), while NfL immunoreactivity showed the opposite, being more abundant in superficial than in deep cortical layers in all groups (controls: +26% compared to deep layers, *p<0.001*; PD: +27%, *p<0.001*; PDD/DLB: +21%, *p=0.003*). For more details on layer-specific distribution of neurofilaments, see Supplementary Fig. 6, online resource 2.

**Fig. 2.**
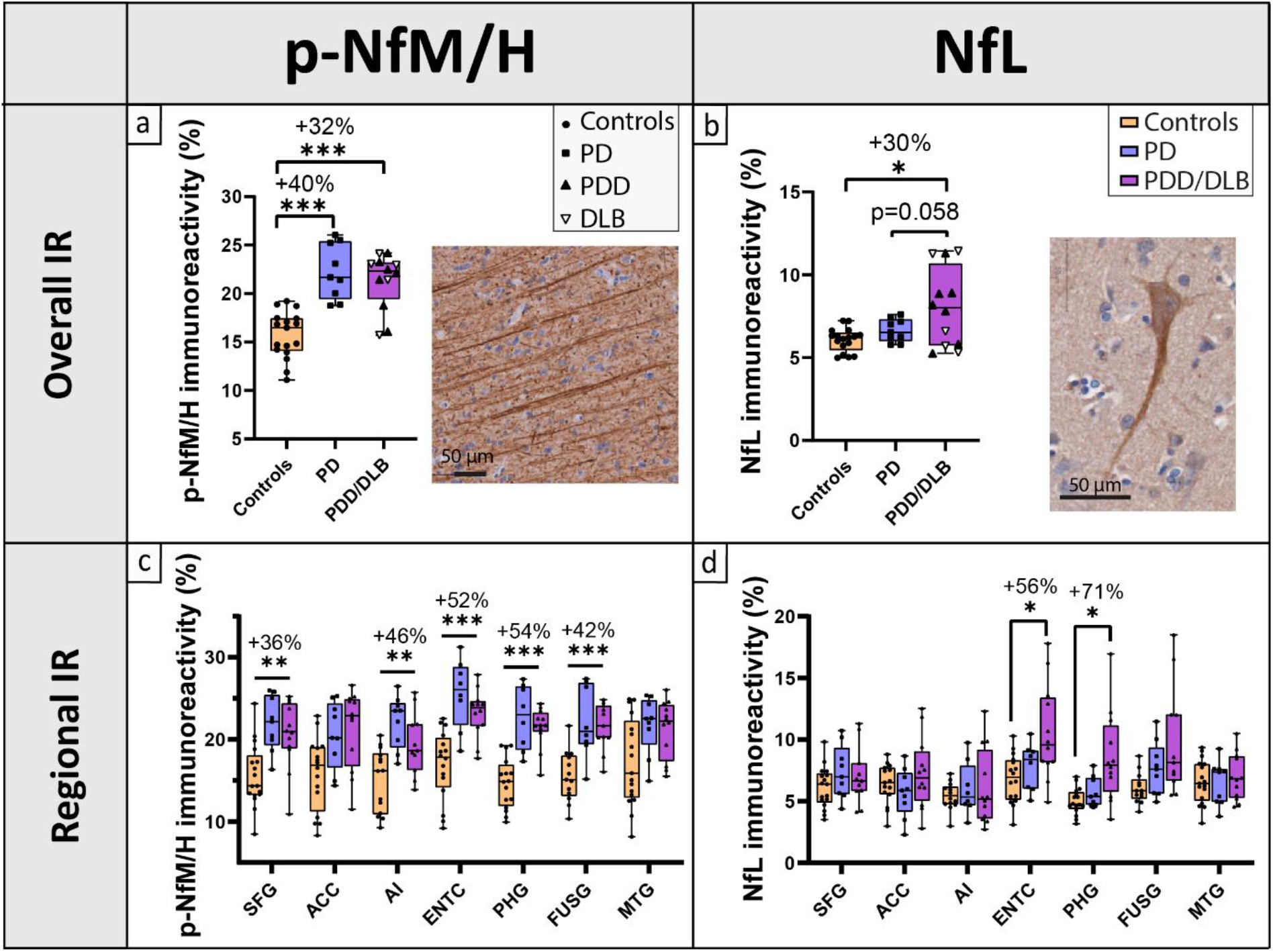
Cortical neurofilament immunoreactivity (IR) and distribution across cortical brain regions. **a** and **c** cortical p-NfM/H and **b** and **d** NfL immunoreactivity are shown for controls, PD and PDD/DLB groups. The first row **a** and **b** shows the overall neurofilament immunoreactivity across all cortical regions examined, with percentage differences compared to controls indicated at the top of the graph; every data point represents a donor averaged across all cortical regions, and the clinical groups are indicated by shapes, showing that PDD and DLB donors behaved similarly. Moreover, images of p-NfM/H **a** and NfL stainings **b** show that p-NfM/H targets the axons, while NfL targets both the neuronal soma and its processes. The second row **c** and **d** shows the regional neurofilament immunoreactivity in the 7 cortical regions analysed. Percentage differences **c** between p-NfM/H immunoreactivity in PD vs controls and **d** between NfL immunoreactivity in PDD/DLB vs controls are shown for brain areas hosting a significant difference. From these data, we can infer that p-NfM/H was uniformly increased across cortical areas analysed in both PD and PDD/DLB, while NfL was increased specifically in the entorhinal and parahippocampal cortex in PDD/DLB. Legend: *\*p<0.05, **p<0.01, ***p<0.001 when compared to controls; ACC: anterior cingulate cortex; AI: anterior insula; DLB: Dementia with Lewy Bodies; ENTC: entorhinal cortex; FUSG: fusiform gyrus; IR: immunoreactivity; MTG: middle temporal gyrus; NfL: neurofilament light chain; PHG: parahippocampal gyrus; PD: Parkinson’s disease; PDD: Parkinson’s disease dementia; p-NfM/H: phosphorylated neurofilament medium and heavy chain; SFG: superior frontal gyrus*

Overall, p-NfM/H immunoreactivity was significantly increased in both PD (*p<0.001*) and PDD/DLB (*p<0.001*) compared to controls by 40% and 32%, respectively (**Fig. 2a** and Supplementary Fig. 3a, online resource 2, for representative images). To investigate whether the increase in p-NfM/H immunoreactivity in the PDD/DLB group was driven by a specific clinical phenotype (PDD or DLB), we explored the difference in p-NfM/H immunoreactivity between these groups, but observed no differences. Regionally, p-NfM/H immunoreactivity was significantly increased in both PD and PDD/DLB groups compared to controls in the superior frontal gyrus (PD: +36%, *p-FDR=0.009;* PDD/DLB: +27%, *p-FDR=0.036*), entorhinal cortex (PD: +52%, *p-FDR=0.001;* PDD/DLB: +38%, *p-FDR=0.003*), parahippocampal gyrus (PD: +54%, *p-FDR<0.001;* PDD/DLB: +46%, *p-FDR<0.001*), fusiform gyrus (PD: +42%, *p-FDR=0.019;* PDD/DLB: +39%, *p-FDR=0.005*) and anterior insula (PD: +46%, *p-FDR=0.003;* PDD/DLB: +29%, *p-FDR=0.049*) (**Fig. 2c**).

Overall, NfL immunoreactivity was significantly increased in the PDD/DLB group compared to controls (+30%, *p=0.037*), and tended to be increased in PDD/DLB compared to PD (*p=0.058*), while it did not differ in PD compared to controls (*p=1.000*) (**Fig. 2b,** and Supplementary Fig. 3b, online resource 2, for representative images). To investigate whether the increase in NfL immunoreactivity in the PDD/DLB group was driven by a specific clinical phenotype (PDD or DLB), we explored the difference in NfL immunoreactivity between these groups, but observed no differences. Regionally, NfL immunoreactivity was increased specifically in the entorhinal cortex (+56%, *p-FDR=0.028*) and the parahippocampal gyrus (+71%, *p-FDR=0.016*) in PDD/DLB compared to controls (**Fig. 2d**).

Taken together, p-NfM/H immunoreactivity was uniformly increased across all cortical areas analysed in all patient groups, while increased NfL immunoreactivity was observed specifically in the entorhinal cortex and the parahippocampal gyrus in PDD/DLB.

### Accumulation and fragmentation of neurofilaments in diseased neurons in PDD/DLB

To better understand the observed increase in p-NfM/H and NfL immunoreactivity in PD and PDD/DLB, we examined the morphological differences of both neurofilaments in the region that was most affected, the parahippocampal gyrus, using multi-label immunofluorescence and 3D CSLM.

First, NfL was much more present in neuronal somas and apical axons of PDD/DLB compared to PD and controls (**Fig. 3a-c**). Specifically, NfL seemed to accumulate in neurons showing neurodegenerative morphologies, such as nuclear fragmentation and swelling (**Fig. 3c**), and ballooning and corkscrew deformation of the soma (**Fig. 3d)**. Second, accumulation and fragmentation of NfL staining patterns were observed in several axons, where axonal swellings intermitted axonal fragmentations in both PD and, more abundantly, in PDD/DLB (**Fig. 3e, g, h,** and **i**), which was not observed in controls. Third, we also observed glial cells close to an apparently fragmented axon in PDD/DLB (**Fig. 3c, e**). Lastly, while p-NfM/H and NfL co-localized in most of control axons, axons in the deep cortical layers of PD and PDD/DLB cases were mostly positive to p-NfM/H or NfL, but rarely the two neurofilaments co-localized (**Fig. 3f-h**). This was also supported by our quantitative data, showing an absence of significant correlation between the two neurofilaments in PD and PDD/DLB groups, which was present in controls (Supplementary Fig. 7, online resource 2).

**Fig. 3.**
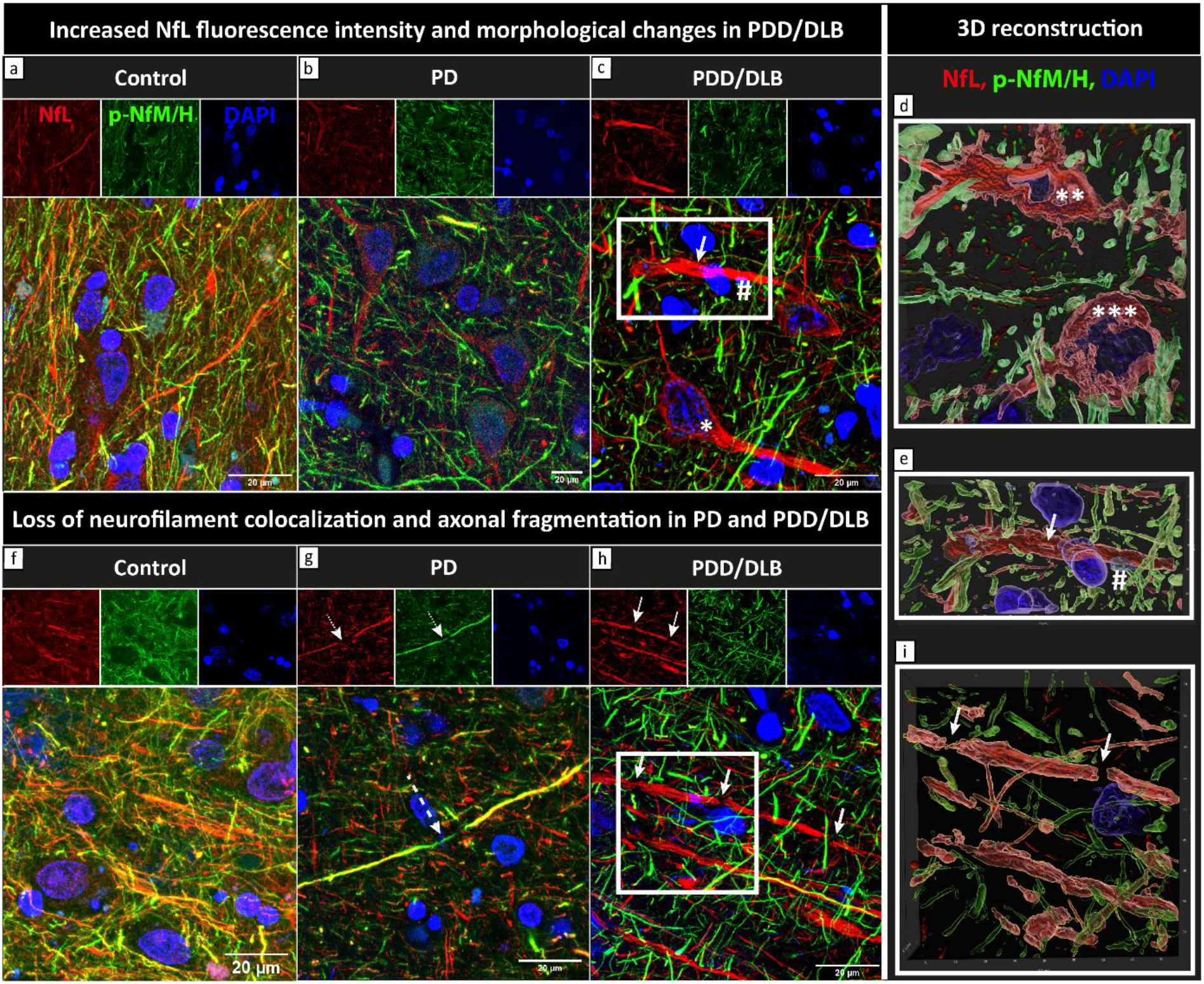
NfL accumulation and fragmentation, and loss of neurofilament colocalization in PD and PDD/DLB. Maximum projections (z-stacks n=6 μm) of fluorescence stainings of NfL (red) and p-NfM/H (green) together with DAPI (blue) in the parahippocampal gyrus of a representative control (*left*, control number 10 in Supplementary Table 1, online resource 1), PD (*middle*, PD number 3 in Supplementary Table 1, online resource 1) and PDD/DLB (*right*, here a DLB donor, PDD/DLB number 12 in Supplementary Table 1, online resource 1) are shown. **a**, **b**, and **c** show the difference in NfL staining pattern in control, PD and PDD/DLB, respectively. NfL was markedly accumulated in neuronal somas and proximal axons in PDD/DLB cases, reflecting the higher NfL immunoreactivity described quantitatively in **Fig. 2d.** High NfL immunoreactivity was found in neuronal somas showing neurodegenerative morphological features, such as **c** neurons with nuclear fragmentation and swelling (*), and **d** corkscrew (**) and ballooned (***) appearances (panel **d** shows a zoom-in 3D surface-reconstruction of neurodegenerative neurons). Moreover, **c** NfL accumulation was also identified in a proximal axon showing fragmentation (arrow), closely tighten by a glial cell (#); **e** shows the zoom-in 3D surface-reconstruction of the white square in **c**, showing axonal fragmented morphology (arrow) and a glial cell wrapping the degenerating axon (#). **f**, **g** and **h** show the colocalization (yellow) of NfL and p-NfM/H in control, PD and PDD/DLB, respectively, which was reduced in PD and PDD/DLB. Moreover, accumulation and fragmentation of NfL staining patterns were observed in several axons, where **g** NfL seemed to fragment within a p-NfM/H axon in PD (dashed arrow) and **h** axonal swellings intermitted axonal fragmentations in PDD/DLB (arrows). **i** shows the zoom-in 3D surface-reconstruction of the white square in **h**, clearly showing fragmentation of the NfL positive axon (arrows). Legend: *DLB: Dementia with Lewy Bodies; NfL: neurofilament light chain; PD: Parkinson’s disease; PDD: Parkinson’s disease dementia; p-NfM/H: phosphorylated neurofilament medium and heavy chain*

Taken together, NfL seemed to accumulate in diseased neurons in PDD/DLB and to fragment axons in PD and, more abundantly, in PDD/DLB. Moreover, both neurofilaments showed less overlap in immunoreactivity profile in PD and PDD/DLB groups.

### Cortical neurofilament immunoreactivity correlates with pSer129-αSyn, and more strongly with p-tau load in PDD/DLB

No correlations were found between p-NfM/H immunoreactivity and pSer129-αSyn pathology in any group across all regions (*p>0.05*, **Fig. 4a**). On the other hand, p-NfM/H positively correlated with p-tau load in the PDD/DLB group across all regions (*r = 0.29, R^2^=8%, p=0.043*), which was a trend in PD (*r = 0.32, p=0.066*), and absent in controls (*p=0.149*) (**Fig. 4b**). No correlation was found with Aβ load (*p>0.05*). The associations were not specific to the superficial or deep cortical layers (for all correlations, see Supplementary Table 5, online resource 1).

**Fig. 4.**
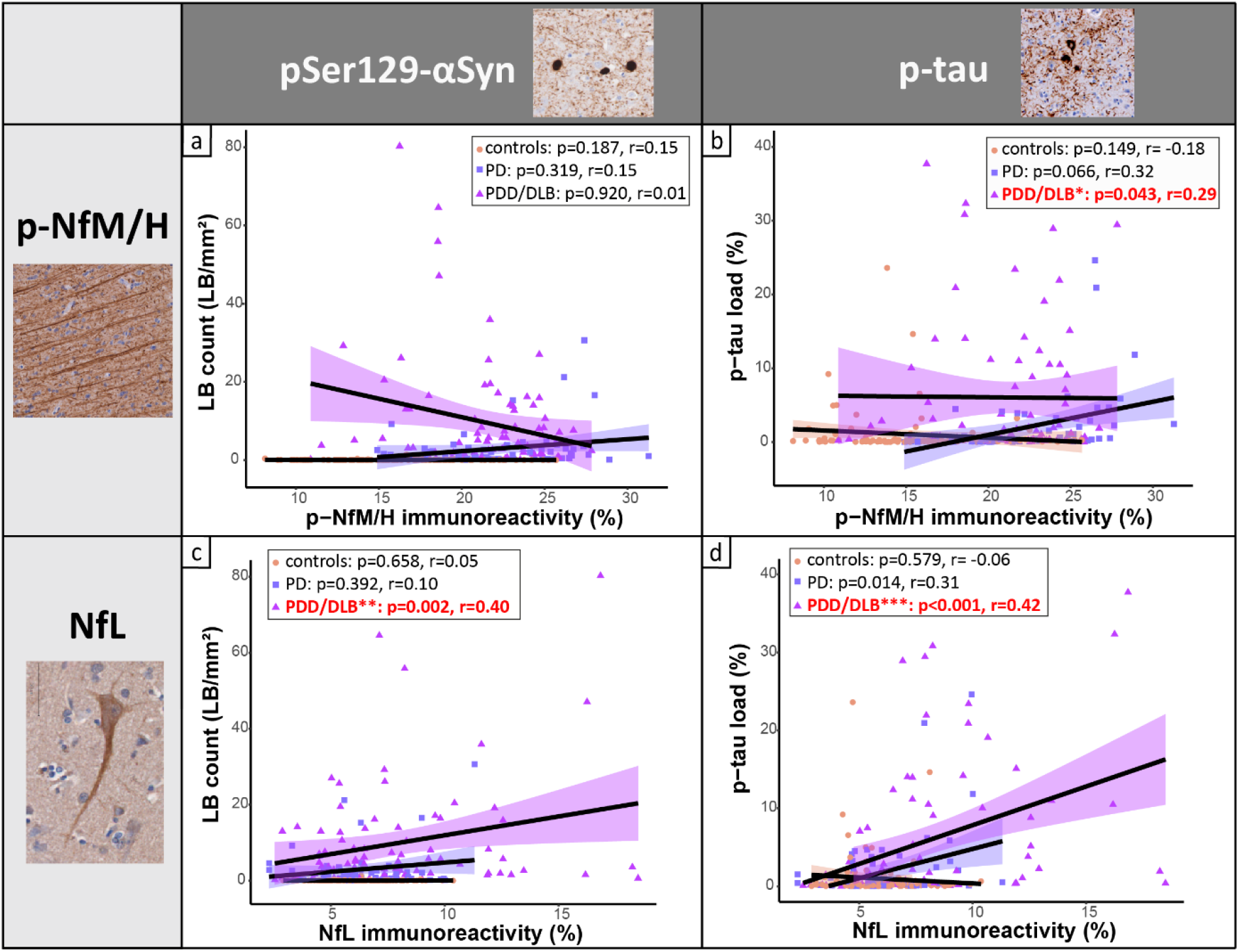
Cortical neurofilament immunoreactivity correlates with pSer129-αSyn and p-tau load. **a** and **b** correlation of **a** and **b** cortical p-NfM/H and **c** and **d** NfL immunoreactivity with pSer129-αSyn-positive LB count and p-tau load, respectively. Every data point represents a cortical region of a donor, and the groups are indicated by colours in **a, b, c** and **d**. For each group, the regression line is shown, together with its colour-coded standard error (orange for controls, purple for PD and pink for PDD/DLB). Significant correlations are highlighted in bold red in the upper box in every panel. Legend: correlation significant at *\*p<0.05, **p<0.01, ***p<0.001; DLB: Dementia with Lewy Bodies; LB: Lewy Body; NfL: neurofilament light chain; PD: Parkinson’s disease; PDD: Parkinson’s disease dementia; p-NfM/H: phosphorylated neurofilament medium and heavy chain; p-tau: phosphorylated tau*

NfL positively correlated with pSer129-αSyn pathology in PDD/DLB across all regions (LB count: *r = 0.40, R^2^=16%, p=0.002;* pSer129-αSyn load: *r = 0.36, R^2^=13%, p=0.005*), but not in the PD group (LB count: *p=0.392;* pSer129-αSyn load: *p=0.910*) (**Fig. 4c**). Moreover, NfL also positively correlated with p-tau load in PDD/DLB (*r = 0.42, R^2^=18%, p<0.001*) and PD (*r = 0.31, R^2^=10%, p=0.014*), but not in controls across all regions (*p=0.579*) (**Fig. 4d**). When both pSer129-αSyn and p-tau load were considered, NfL correlated with p-tau load (*r = 0.24, R^2^=6%, p=0.030*) but not pSer129-αSyn load (*p=0.195*). Taken together, it seems that NfL correlated more strongly with p-tau than pSer129-αSyn load. No correlation was found between NfL and Aβ load (*p>0.05*). Additionally, the associations were not specific to superficial or deep cortical layers (for all correlations, see Supplementary Table 6, online resource 1).

Taken together, in PDD/DLB, higher p-NfM/H immunoreactivity correlated with higher p-tau load, while NfL immunoreactivity correlated with higher pSer129-αSyn load, but more strongly with higher p-tau load. None of the neurofilaments correlated with Aβ load.

### Cortical neurofilament immunoreactivity is reflected by MRI cortical thickness in PD

To investigate whether MRI-derived cortical thickness is sensitive to regional neurofilament changes in the cortex of PD and PDD/DLB donors, we assessed whether regional neurofilament immunoreactivity correlated with cortical thickness across the included brain regions.

On MRI, neither PD (*p=1.000*) or PDD/DLB cases (*p=0.756*) showed cortical thickness differences compared to controls across and within all cortical brain regions (all *p-FDR>0.05*) (Supplementary Fig. 8a and b, online resource 2). Moreover, cortical thickness did not correlate with any pathological hallmark (*p>0.05*) (Supplementary Fig. 9a and c, online resource 2, and Supplementary Table 7, online resource 1).

P-NfM/H immunoreactivity negatively correlated with cortical thickness in the full PD+PDD/DLB cohort across all cortical regions (*r = −0.23, R^2^=5%, p=0.020*). Particularly, the association was driven by the PD group (*r = −0.32, R^2^=10%, p=0.047*), and not significant in PDD/DLB (*r = −0.21, p=0.106*) (**Fig. 5a**). No correlations were found in the control group (*p>0.05*), or for individual brain regions (*p-FDR>0.05*).

**Fig. 5.**
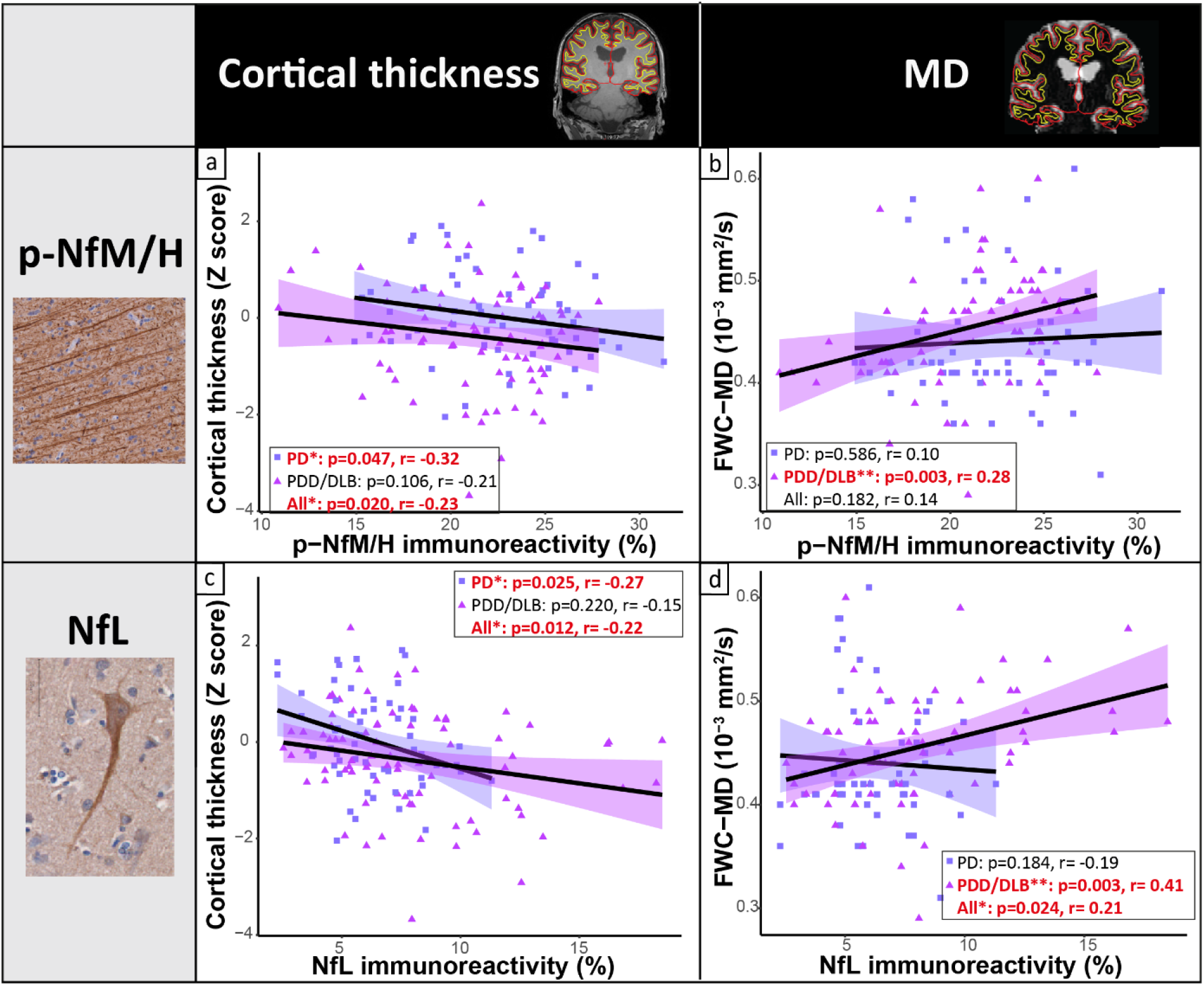
Cortical neurofilament immunoreactivity correlates with decreased cortical thickness in PD and increased cortical MD in PDD/DLB. **a** and **b** correlation of cortical p-NfM/H and **c** and **d** NfL immunoreactivity with cortical thickness on the left, and cortical MD on the right. Both neurofilaments immunoreactivity correlated with decreased cortical thickness, specifically in PD, and with higher cortical MD, specifically in PDD/DLB. Every data point represents a cortical region of interest of a donor, and the groups are indicated by colours in the figure. For each group, the regression line is shown, together with its colour-coded standard error (purple for PD and pink for PDD/DLB). Significant correlations are highlighted in red bold in the box in every panel. Legend: correlation significant at *\*p<0.05, **p<0.01, ***p<0.001; DLB: Dementia with Lewy Bodies; FWC-MD: free-water corrected mean diffusivity; MD: mean diffusivity; NfL: neurofilament light chain; PD: Parkinson’s disease; PDD: Parkinson’s disease dementia; p-NfM/H: phosphorylated neurofilament medium and heavy chain*

Additionally, NfL immunoreactivity negatively correlated with cortical thickness in the full PD+PDD/DLB cohort across all cortical regions (*r = −0.22, R^2^=5%, p=0.012*). Again, the association was driven by PD (*r = −0.27, R^2^=7%, p=0.025*), and absent in PDD/DLB (*r = −0.15, p=0.220*) (**Fig. 5c**). No correlations were found in the control group (*p>0.05*), or for individual brain regions (*p-FDR>0.05*).

Taken together, cortical thickness is not sensitive to pathological load. However, both p-NfM/H and NfL immunoreactivity levels correlated with cortical thickness, especially in the PD group.

### Cortical neurofilaments immunoreactivity is reflected by cortical diffusivity in PDD/DLB

To investigate whether cortical MD is sensitive to regional neurofilament variation in the cortex of PD and PDD/DLB donors, we assessed whether regional neurofilament immunoreactivity correlated with MD across the cortical regions of interest.

Neither PD (*p=1.000*) or PDD/DLB donors (*p=0.487*) showed overall cortical MD differences compared to controls across and within the regions analysed (*p-FDR>0.05*) (Supplementary Fig. 8c and d, online resource 2). Overall, cortical MD positively correlated with pSer129-αSyn (*r = 0.32, R^2^=10%, p=0.002*) and p-tau pathology (*r = 0.36, R^2^=13%, p<0.001*) in PDD/DLB across all cortical regions, but not with Aβ load (*p=0.339*) (Supplementary Fig. 9b and d, online resource 2, and Supplementary Table 8, online resource 1).

P-NfM/H immunoreactivity did not correlated with cortical MD in the full PD+PDD/DLB cohort across all cortical regions (*p=0.182*). However, p-NfM/H immunoreactivity positively correlated with cortical MD in the PDD/DLB group (*r = 0.28, R^2^=8%, p=0.003*) (**Fig. 5b**). No correlations were found in the control group (*p=0.430*), PD group (*p=0.586*), or for individual brain regions (*p-FDR>0.05*).

Additionally, NfL immunoreactivity positively correlated with cortical MD in the full PD+PDD/DLB cohort across all cortical regions (*r = 0.21, R^2^=4%, p=0.024*). Particularly, the association was present in the PDD/DLB group (*r = 0.41, R^2^=17%, p=0.003*), but not in the PD (*p=0.184*) or control group (*p=0.738*) (**Fig. 5d**). No correlations were found for individual brain regions (*p-FDR>0.05*).

Taken together, cortical MD is a sensitive marker for pSer129-αSyn and p-tau load. In addition, both p-NfM/H and NfL immunoreactivity correlated with increased cortical MD, especially in the PDD/DLB group.

## Discussion

Using a post-mortem within-subject *in-situ* MRI and histopathology approach [40], we investigated cortical regional neurofilament immunoreactivity and its relation to pathological hallmarks and MRI outcome measures in PD and PDD/DLB from a microscale to a mesoscale level. We found that p-NfM/H was increased in PD and PDD/DLB across almost all cortical regions, while NfL was increased specifically in the parahippocampal gyrus and entorhinal cortex of PDD/DLB donors. NfL accumulated in somas of diseased neurons showing neurodegenerative morphologies, and in axons showing fragmentation. Furthermore, we found that in PDD/DLB, p-NfM/H immunoreactivity levels positively correlated with p-tau load, whereas NfL positively correlated with pSer129-αSyn pathology and more strongly with p-tau load. Lastly, we showed that neurofilament immunoreactivity correlated with cortical thinning in PD and with higher cortical MD in PDD/DLB.

For p-NfM/H, we found that both PD and PDD/DLB donors had higher p-NfM/H immunoreactivity levels compared to controls, and that this increase was uniform across the analysed cortical areas. An increase in phosphorylation of neurofilament medium and heavy chains (NfM/H) has been described in several neurological disorders, such as AD, multiple sclerosis, amyotrophic lateral sclerosis and stroke [55], but it has never been investigated in the cerebral cortex of PD patients. Phosphorylation is assumed to be the dominant phosoform of both NfM and NfH, and under normal conditions axons show extensive neurofilament phosphorylation, while there is little or no phosphorylation in the neuronal soma and its dendrites [32, 55], which we also show in our study. Neurofilament phosphorylation is a highly regulated and complex process, and an increased phosphorylation of subunits M (medium) and H (heavy), as found in our study, has been linked to several pathological processes [32, 55]. Firstly, neurofilament hyper-phosphorylation seems to impair axonal transport, which has been linked to a stoichiometric imbalance of neurofilament subunits [32, 55]. Interestingly, we found an absence of colocalization (qualitatively) and correlation (quantitatively) between p-NfM/H and NfL immunoreactivity in PD and PDD/DLB, which resembles a stoichiometric imbalance in neurofilament subunits. Secondly, neurofilament hyper-phosphorylation might be a by-product of dysregulated kinase activity due to stress factors, such as chronic oxidant stress in neurodegenerative diseases [32]. Cellular stress factors are associated with extensive phosphorylation of cytoskeletal elements through several proline-directed kinases, which are capable of phosphorylating neurofilaments and tau [32, 43], and have been shown to be dysregulated in AD and PD [17, 43]. Once phosphorylated, NfM/H promotes neurofilament-neurofilament interactions, leading to neurofilament accumulation [32, 43], which we also found in our study. Moreover, we found that both p-NfM/H and NfL correlated to p-tau load in PDD/DLB, suggestive of cytoskeletal alterations in axons in cortical brain regions [32, 43]. Taken together (**Fig. 6)**, increased NfM/H phosphorylation across the cortex seems to point towards evidence of cellular stress and axonal injury in our PD(D)/DLB cohort, even when donors were not demented, and p-tau pathology was not present. This suggests that PD pathophysiology, independently of AD co-pathology, can also trigger an increase in NfM/H phosphorylation, and therefore axonal stress and injury.

**Fig. 6.**
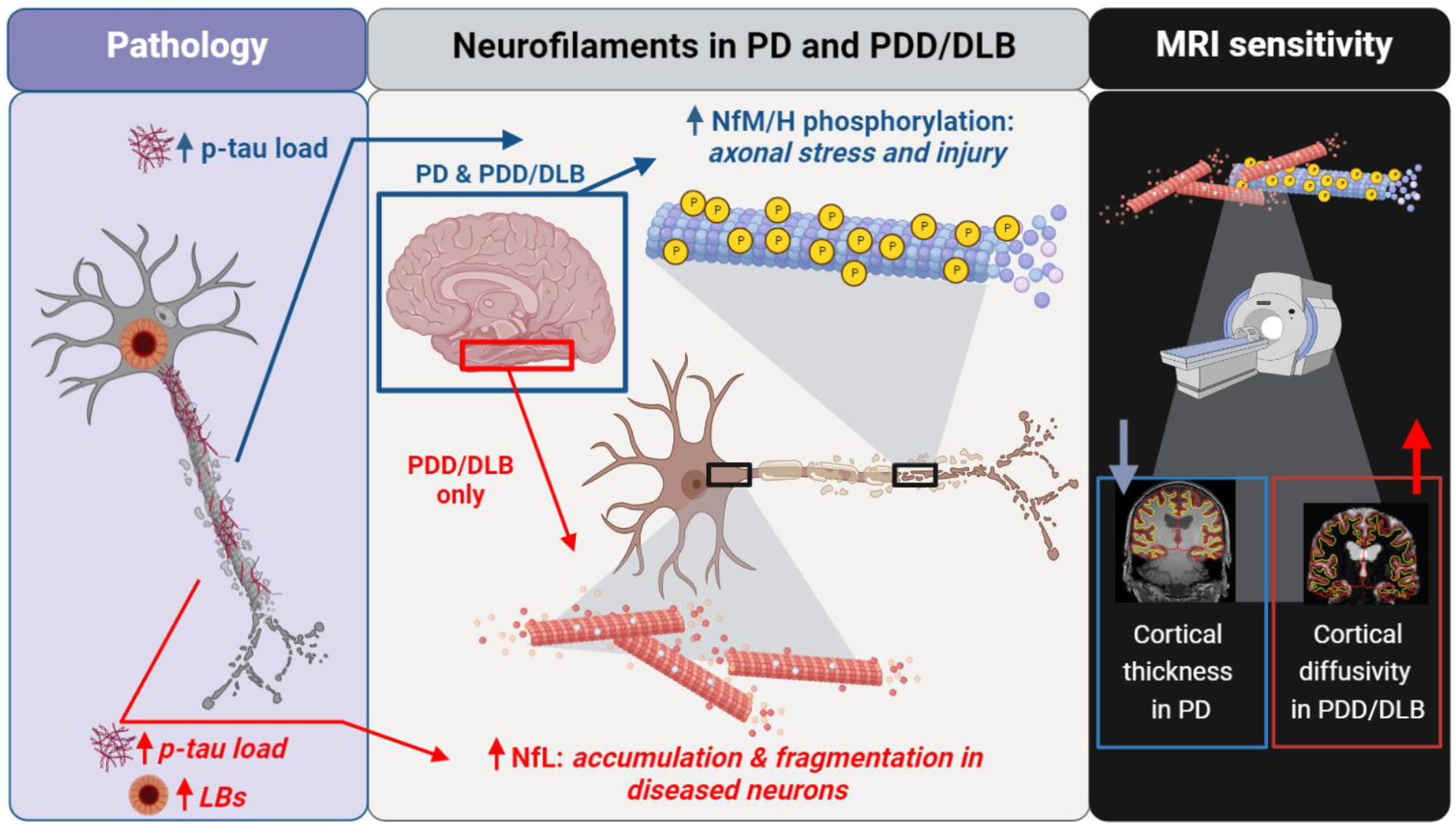
Cortical neurofilaments are increased in PD and PDD/DLB, and associate with pathology load and decreased cortical thickness and increased intra-cortical diffusion on MRI. *Middle panel*: neurofilament immunoreactivity was increased in both PD and PDD/DLB cortex. Specifically, increased NfM/H phosphorylation occurred across the cortex in both PD and PDD/DLB (blue square), indicating axonal stress and injury, and associated with increased p-tau load (*left panel*, blue arrow). On the other hand, an increased NfL immunoreactivity, representing NfL accumulation in disease neurons and fragmented axons, was observed in the entorhinal cortex and parahippocampal gyrus of PDD/DLB donors (red square) and associated with LB count and to p-tau load (*left panel*, red arrow), suggesting structural changes with increasing pathological burden, and confirming its important role in the context of cognitive decline. *Right panel*: both increased p-NfM/H (i.e. axonal stress and injury) and NfL immunoreactivity (i.e. NfL accumulation and fragmentation) were reflected by cortical thinning in PD and increased cortical MD in PDD/DLB. Figure created with BioRender.com. Legend: *DLB: Dementia with Lewy Bodies; LB: Lewy Body; NfL: neurofilament light chain; PD: Parkinson’s disease; PDD: Parkinson’s disease dementia; NfM/H: phosphorylated neurofilament medium and heavy chain; p-tau: phosphorylated tau*

Furthermore, we found that PDD/DLB cases had higher cortical NfL immunoreactivity levels compared to age-matched controls, and that this increase was specific to the entorhinal cortex and parahippocampal gyrus. An increase in NfL immunoreactivity seems counterintuitive, as it is generally believed that breakdown of axons leads to an increase of this marker in the extracellular fluid, and therefore a decrease of it in brain tissue [41]. However, higher NfL immunoreactivity has been described before in an animal model and human brain tissue of stroke patients [30, 48]. Specifically, it has been shown that the same NfL antibody used in this study, the polyclonal rabbit NfL (AA 1-284), not only targets NfL, but most importantly also targets its degradation products [48]. Translating this information to our study, we recorded an increase of NfL immunoreactivity in PDD/DLB, which may result from increased NfL degradation products. In support of this, we also found that NfL accumulated in neuronal somas that showed patterns of neurodegeneration (e.g. nuclear material fragmentation, ballooned and corkscrew cell bodies), and NfL positive axons often showed fragmentation, as described before in a mouse model of stroke [30, 48]. Taken together, increased NfL in para-hippocampal regions seems to point towards evidence of neurofilament accumulation and fragmentation in PDD and DLB. These results are consistent with several CSF and plasma studies, which report that higher CSF or plasma NfL levels correlate with cognitive decline in PD [7, 9, 44, 45, 67]. The information that we add to these studies is regional, by showing that NfL immunoreactivity is specifically increased in the entorhinal and parahippocampal cortex, regions that are strongly involved in cognitive processes [66], and are the first to show p-tau accumulation in the adult and aged human brain [14]. This process might therefore hint towards why NfL CSF and plasma levels are increased in several neurological disorders which are per definition p-tau positive, such as AD, frontotemporal dementia and multiple system atrophy [7, 41]. Moreover, beside NfL correlation with p-tau load, we showed that NfL immunoreactivity levels also associated with pSer129-αSyn load. CSF NfL levels were shown to positively correlate with CSF αSyn [28] and with decline in memory, attentional and executive functioning in PD [53]. Additionally, we recently showed that neurofilaments accumulate and cluster around the core of Lewy bodies in post-mortem brain tissue of PD(D)/DLB donors [52], illustrating a close relationship between NfL and pathological αSyn, suggesting a role for neurofilament in encapsulating toxic proteins. Taken together (**Fig. 6)**, aggregation and fragmentation of NfL, and its association to p-tau and pSer129-αSyn pathology, suggest that NfL undergoes structural changes with increasing pathological burden, confirming its important role in the context of cognitive decline.

To investigate the possibility of measuring the above-mentioned pathological changes in clinical practice, we evaluated whether cortical pathology and neurofilaments immunoreactivity would be reflected by MRI biomarkers of neurodegeneration, including cortical thickness and cortical microstructural integrity. We found that increased neurofilament immunoreactivity correlated with lower cortical thickness in PD, and with higher cortical MD in PDD/DLB. This discrepant association in non-demented and demented cases may be explained by the fact that we found cortical MD to be a more sensitive marker than cortical thickness in picking up pSer129-αSyn and p-tau pathology, which was more abundantly present in PDD/DLB than in non-demented PD. Our results partially go in line with previous literature, where serum NfL was shown to correlate with cortical thinning of several brain areas in *de-novo* PD, but also with increased cortical MD [59]. Unfortunately, no similar studies have been carried out in PDD and DLB, making this the first study to investigate this relationship. Taken together (**Fig. 6)**, we found that cortical thinning in PD, and lower microstructural integrity in PDD/DLB seem to reflect not only accumulation of neurofilaments (i.e. increased NfL), but also cellular stress and axonal damage (i.e. increased p-NfH/M).

The main strength of this study is the combination of MRI and gold-standard histological data of multiple cortical regions, which were collected from well-characterized donors. All donors had pathological confirmation of clinical diagnosis, as clinical-pathological discrepancies occur in about 15% of cases and might obscure *in-vivo* studies [26, 34]. We studied both quantitatively and qualitatively regional cortical neurofilament changes in PD. NfL is widely used as a CSF or blood biomarker to detect axonal degeneration in patient cohorts, and with this study we hope to shed light on the regional and morphological changes in neurofilaments, linking neurodegenerative features from micro-to mesoscale and to the clinic. Therefore, results of this study provide evidence to the important roles of neurofilaments in neurodegeneration and their MRI signatures.

There are also some limitations. Neurofilament changes are not a process specific to PD(D) or DLB. In fact, increased CSF and plasma NfL levels have been described in several other neurological disorders compared to age-matched controls, and are considered a measure of neuroaxonal damage independent of casual pathways [7, 41]. Moreover, neurofilament changes only account for part of the variance of the damage seen on MRI and DTI (in our study up to 17%), and it is likely that other cellular and molecular components contribute to neurodegeneration, such as synaptic loss or glial changes. In addition, we did not find associations between MRI outcome measures and neurofilament immunoreactivity within the included brain regions (only across). This lack of regional sensitivity to neurofilament changes might be due to our small sample size, since large PD(D) cohorts are needed to pick up subtle changes in cortical thinning [42]. Future research should therefore investigate the mechanisms behind the increase in neurofilaments and their association to MRI biomarkers in larger cohorts and in other neurological disorders showing high NfL levels and more pronounced regional atrophy, such as AD, frontotemporal dementia and multiple system atrophy [7, 41].

Taken together, we provide evidence for increased phosphorylation of NfM/H, suggesting axonal stress and injury across the cortex of PD, PDD and DLB donors. Furthermore, we show that NfL immunoreactivity was increased in the parahippocampal gyrus and entorhinal cortex of PDD and DLB donors, reflecting aggregation and fragmentation of NfL in diseased neurons and axons, suggesting that NfL undergoes structural changes with increasing pathological burden in the context of cognitive decline. Importantly, we provide evidence for the relation between pathological burden and neurofilament levels, and demonstrate that such microscopic markers at least partly explain MRI markers that are associated with the neurodegenerative process.

## Abbreviations

Aβ: amyloid-beta
AD: Alzheimer’s disease
CSF: cerebrospinal fluid
DLB: dementia with Lewy bodies
DTI: diffusion tensor imaging
FDR: false discovery rate
FWC-MD: free-water corrected mean diffusivity
IHC: immunohistochemistry
LB: Lewy body
MD: mean diffusivity
NfL: neurofilament light chain
NFT: neurofibrillary tangles
PD: Parkinson’s disease
PDD: Parkinson’s disease dementia
p-NfM/H: phosphorylated neurofilament medium and heavy chain
p-tau: phosphorylated-tau
pSer129-αSyn: phosphorylated Ser129 alpha synuclein

## Supplementary information

Supplementary Information The online version contains supplementary material available as open resource material, and includes Supplementary Tables (open resource 1), Supplementary Figures (open resource 2), and Supplementary Material (open resource 3).

## Acknowledgements

We would like to thank all brain donors and their next of kin for brain donation. We would also like to thank the autopsy teams of the Netherlands Brain Bank (NBB) and Normal Aging Brain collection Amsterdam (NABCA).

## References

1 Aarsland D, Zaccai J, Brayne C (2005) A systematic review of prevalence studies of dementia in Parkinson’s disease. Movement disorders: official journal of the Movement Disorder Society 20: 1255–1263

2 Adler DH, Pluta J, Kadivar S, Craige C, Gee JC, Avants BB, Yushkevich PA (2014) Histology-derived volumetric annotation of the human hippocampal subfields in postmortem MRI. Neuroimage 84: 505–523

3 Alafuzoff I, Arzberger T, Al-Sarraj S, Bodi I, Bogdanovic N, Braak H, Bugiani O, Del-Tredici K, Ferrer I, Gelpi E (2008) Staging of neurofibrillary pathology in Alzheimer’s disease: a study of the BrainNet Europe Consortium. Brain pathology 18: 484–496

4 Alafuzoff I, Ince PG, Arzberger T, Al-Sarraj S, Bell J, Bodi I, Bogdanovic N, Bugiani O, Ferrer I, Gelpi E (2009) Staging/typing of Lewy body related α-synuclein pathology: a study of the BrainNet Europe Consortium. Acta neuropathologica 117: 635–652

5 Alafuzoff I, Thal DR, Arzberger T, Bogdanovic N, Al-Sarraj S, Bodi I, Boluda S, Bugiani O, Duyckaerts C, Gelpi E (2009) Assessment of β-amyloid deposits in human brain: a study of the BrainNet Europe Consortium. Acta neuropathologica 117: 309–320

6 Arendt T, Morawski M, Gartner U, Frohlich N, Schulze F, Wohmann N, Jager C, Eisenloffel C, Gertz HJ, Mueller W et al (2017) Inhomogeneous distribution of Alzheimer pathology along the isocortical relief. Are cortical convolutions an Achilles heel of evolution? Brain Pathol 27: 603–611 Doi 10.1111/bpa.12442

7 Ashton NJ, Janelidze S, Al Khleifat A, Leuzy A, van der Ende EL, Karikari TK, Benedet AL, Pascoal TA, Lleó A, Parnetti L (2021) A multicentre validation study of the diagnostic value of plasma neurofilament light. Nature communications 12: 1–12

8 Atkinson-Clement C, Pinto S, Eusebio A, Coulon O (2017) Diffusion tensor imaging in Parkinson’s disease: review and meta-analysis. Neuroimage: Clinical 16: 98–110

9 Bäckström D, Linder J, Mo SJ, Riklund K, Zetterberg H, Blennow K, Forsgren L, Lenfeldt N (2020) NfL as a biomarker for neurodegeneration and survival in Parkinson disease. Neurology 95: e827–e838

10 Bankhead P, Loughrey MB, Fernandez JA, Dombrowski Y, McArt DG, Dunne PD, McQuaid S, Gray RT, Murray LJ, Coleman HG et al (2017) QuPath: Open source software for digital pathology image analysis. Sci Rep 7: 16878 Doi 10.1038/s41598-017-17204-5

11 Benjamini Y, Hochberg Y (1995) Controlling the false discovery rate: a practical and powerful approach to multiple testing. Journal of the Royal statistical society: series B (Methodological) 57: 289–300

12 Beyer MK, Larsen JP, Aarsland D (2007) Gray matter atrophy in Parkinson disease with dementia and dementia with Lewy bodies. Neurology 69: 747–754

13 Boon BDC, Pouwels PJW, Jonkman LE, Keijzer MJ, Preziosa P, van de Berg WDJ, Geurts JJG, Scheltens P, Barkhof F, Rozemuller AJM et al (2019) Can post-mortem MRI be used as a proxy for in vivo? A case study. Brain Commun 1: fcz030 Doi 10.1093/braincomms/fcz030

14 Braak H, Alafuzoff I, Arzberger T, Kretzschmar H, Del Tredici K (2006) Staging of Alzheimer disease-associated neurofibrillary pathology using paraffin sections and immunocytochemistry. Acta neuropathologica 112: 389–404

15 Braak H, Del Tredici K, Rub U, de Vos RA, Jansen Steur EN, Braak E (2003) Staging of brain pathology related to sporadic Parkinson’s disease. Neurobiol Aging 24: 197–211 Doi 10.1016/s0197-4580(02)00065-9

16 Burton EJ, McKeith IG, Burn DJ, Williams ED, O’Brien JT (2004) Cerebral atrophy in Parkinson’s disease with and without dementia: a comparison with Alzheimer’s disease, dementia with Lewy bodies and controls. Brain 127: 791–800

17 Cheung ZH, Ip NY (2012) Cdk5: a multifaceted kinase in neurodegenerative diseases. Trends in cell biology 22: 169–175

18 Chiapponi C, Piras F, Piras F, Fagioli S, Caltagirone C, Spalletta G (2013) Cortical grey matter and subcortical white matter brain microstructural changes in schizophrenia are localised and age independent: a case-control diffusion tensor imaging study. PLoS One 8: e75115 Doi 10.1371/journal.pone.0075115

19 Cicchetti DV (1994) Guidelines, criteria, and rules of thumb for evaluating normed and standardized assessment instruments in psychology. Psychological assessment 6: 284

20 Daducci A, Gerhard S, Griffa A, Lemkaddem A, Cammoun L, Gigandet X, Meuli R, Hagmann P, Thiran JP (2012) The connectome mapper: an open-source processing pipeline to map connectomes with MRI. PLoS One 7: e48121 Doi 10.1371/journal.pone.0048121

21 Dale AM, Fischl B, Sereno MI (1999) Cortical surface-based analysis: I. Segmentation and surface reconstruction. Neuroimage 9: 179–194

22 e Sousa CS, Alarcão J, Martins IP, Ferreira JJ (2022) Frequency of dementia in Parkinson’s disease: A systematic review and meta-analysis. Journal of the Neurological Sciences 432: 120077

23 Emre M, Aarsland D, Brown R, Burn DJ, Duyckaerts C, Mizuno Y, Broe GA, Cummings J, Dickson DW, Gauthier S (2007) Clinical diagnostic criteria for dementia associated with Parkinson’s disease. Movement disorders: official journal of the Movement Disorder Society 22: 1689–1707

24 Frigerio I, Boon BD, Lin C-P, Galis-de Graaf Y, Bol J, Preziosa P, Twisk J, Barkhof F, Hoozemans JJ, Bouwman FH (2021) Amyloid-β, p-tau and reactive microglia are pathological correlates of MRI cortical atrophy in Alzheimer’s disease. Brain communications 3: fcab281

25 Garyfallidis E, Brett M, Amirbekian B, Rokem A, Van Der Walt S, Descoteaux M, Nimmo-Smith I, Contributors D (2014) Dipy, a library for the analysis of diffusion MRI data. Frontiers in neuroinformatics 8: 8

26 Geut H, Hepp D, Foncke E, Berendse H, Rozemuller J, Huitinga I, Van De Berg W (2020) Neuropathological correlates of parkinsonian disorders in a large Dutch autopsy series. Acta neuropathologica communications 8: 1–14

27 Hagmann P, Cammoun L, Gigandet X, Meuli R, Honey CJ, Wedeen VJ, Sporns O (2008) Mapping the structural core of human cerebral cortex. PLoS Biol 6: e159 Doi 10.1371/journal.pbio.0060159

28 Hall S, Surova Y, Öhrfelt A, Zetterberg H, Lindqvist D, Hansson O (2015) CSF biomarkers and clinical progression of Parkinson disease. Neurology 84: 57–63

29 Halliday GM, Song YJC, Harding AJ (2011) Striatal β-amyloid in dementia with Lewy bodies but not Parkinson’s disease. Journal of neural transmission 118: 713–719

30 Härtig W, Krueger M, Hofmann S, Preißler H, Märkel M, Frydrychowicz C, Mueller WC, Bechmann I, Michalski D (2016) Up-regulation of neurofilament light chains is associated with diminished immunoreactivities for MAP2 and tau after ischemic stroke in rodents and in a human case. Journal of Chemical Neuroanatomy 78: 140–148

31 Hepp DH, Vergoossen DL, Huisman E, Lemstra AW, Bank NB, Berendse HW, Rozemuller AJ, Foncke EM, van de Berg WD (2016) Distribution and load of amyloid-β pathology in Parkinson disease and dementia with Lewy bodies. Journal of Neuropathology & Experimental Neurology 75: 936–945

32 Holmgren A, Bouhy D, Timmerman V (2012) Neurofilament phosphorylation and their proline-directed kinases in health and disease. Journal of the Peripheral Nervous System 17: 365–376

33 Hoy AR, Koay CG, Kecskemeti SR, Alexander AL (2014) Optimization of a free water elimination two-compartment model for diffusion tensor imaging. Neuroimage 103: 323–333

34 Hughes AJ, Daniel SE, Ben-Shlomo Y, Lees AJ (2002) The accuracy of diagnosis of parkinsonian syndromes in a specialist movement disorder service. Brain 125: 861–870

35 Insausti R, Munoz-Lopez M, Insausti AM, Artacho-Perula E (2017) The Human Periallocortex: Layer Pattern in Presubiculum, Parasubiculum and Entorhinal Cortex. A Review. Front Neuroanat 11: 84 Doi 10.3389/fnana.2017.00084

36 Irwin DJ, White MT, Toledo JB, Xie SX, Robinson JL, Van Deerlin V, Lee VMY, Leverenz JB, Montine TJ, Duda JE (2012) Neuropathologic substrates of Parkinson disease dementia. Annals of neurology 72: 587–598

37 Jellinger KA (2022) Morphological basis of Parkinson disease-associated cognitive impairment: an update. Journal of Neural Transmission: 1–23

38 Jellinger KA (2003) Neuropathological spectrum of synucleinopathies. Movement disorders: official journal of the Movement Disorder Society 18: 2–12

39 Jenkinson M, Beckmann CF, Behrens TE, Woolrich MW, Smith SM (2012) Fsl. Neuroimage 62: 782–790 Doi 10.1016/j.neuroimage.2011.09.015

40 Jonkman LE, Galis-de Graaf Y, Bulk M, Kaaij E, Pouwels PJ, Barkhof F, Rozemuller AJ, van der Weerd L, Geurts JJ, van de Berg WD (2019) Normal Aging Brain Collection Amsterdam (NABCA): A comprehensive collection of postmortem high-field imaging, neuropathological and morphometric datasets of non-neurological controls. NeuroImage: Clinical 22: 101698

41 Khalil M, Teunissen CE, Otto M, Piehl F, Sormani MP, Gattringer T, Barro C, Kappos L, Comabella M, Fazekas F (2018) Neurofilaments as biomarkers in neurological disorders. Nature Reviews Neurology 14: 577–589

42 Laansma MA, Bright JK, Al-Bachari S, Anderson TJ, Ard T, Assogna F, Baquero KA, Berendse HW, Blair J, Cendes F (2021) International multicenter analysis of brain structure across clinical stages of Parkinson’s disease. Movement disorders 36: 2583–2594

43 Lee H-g, Perry G, Moreira PI, Garrett MR, Liu Q, Zhu X, Takeda A, Nunomura A, Smith MA (2005) Tau phosphorylation in Alzheimer’s disease: pathogen or protector? Trends in molecular medicine 11: 164–169

44 Lerche S, Wurster I, Röben B, Zimmermann M, Machetanz G, Wiethoff S, Dehnert M, Rietschel L, Riebenbauer B, Deuschle C (2020) CSF NFL in a longitudinally assessed PD cohort: age effects and cognitive trajectories. Movement Disorders 35: 1138–1144

45 Lin C-H, Li C-H, Yang K-C, Lin F-J, Wu C-C, Chieh J-J, Chiu M-J (2019) Blood NfL: a biomarker for disease severity and progression in Parkinson disease. Neurology 93: e1104–e1111

46 Lin C-P, Frigerio I, Boon BD, Zhou Z, Rozemuller AJ, Bouwman F, Schoonheim MM, van de Berg WD, Jonkman L (2021) Structural (dys) connectivity associates with cholinergic cell density of the nucleus basalis of Meynert in Alzheimer’s disease. bioRxiv:

47 Lippa C, Duda J, Grossman M, Hurtig H, Aarsland D, Boeve B, Brooks D, Dickson D, Dubois B, Emre M (2007) DLB and PDD boundary issues: diagnosis, treatment, molecular pathology, and biomarkers. Neurology 68: 812–819

48 Mages B, Aleithe S, Altmann S, Blietz A, Nitzsche B, Barthel H, Horn AK, Hobusch C, Härtig W, Krueger M (2018) Impaired neurofilament integrity and neuronal morphology in different models of focal cerebral ischemia and human stroke tissue. Frontiers in Cellular Neuroscience 12: 161

49 McKeith IG, Boeve BF, Dickson DW, Halliday G, Taylor J-P, Weintraub D, Aarsland D, Galvin J, Attems J, Ballard CG (2017) Diagnosis and management of dementia with Lewy bodies: Fourth consensus report of the DLB Consortium. Neurology 89: 88–100

50 Mollenhauer B, Dakna M, Kruse N, Galasko D, Foroud T, Zetterberg H, Schade S, Gera RG, Wang W, Gao F (2020) Validation of serum neurofilament light chain as a biomarker of Parkinson’s disease progression. Movement Disorders 35: 1999–2008

51 Montine TJ, Phelps CH, Beach TG, Bigio EH, Cairns NJ, Dickson DW, Duyckaerts C, Frosch MP, Masliah E, Mirra SS (2012) National Institute on Aging–Alzheimer’s Association guidelines for the neuropathologic assessment of Alzheimer’s disease: a practical approach. Acta neuropathologica 123: 1–11

52 Moors TE, Maat CA, Niedieker D, Mona D, Petersen D, Timmermans-Huisman E, Kole J, El-Mashtoly SF, Spycher L, Zago W (2021) The subcellular arrangement of alpha-synuclein proteoforms in the Parkinson’s disease brain as revealed by multicolor STED microscopy. Acta neuropathologica 142: 423–448

53 Oosterveld LP, Kuiper TI, Majbour NK, Verberk IM, van Dijk KD, Twisk JW, El-Agnaf OM, Teunissen CE, Weinstein HC, Klein M (2020) CSF biomarkers reflecting protein pathology and axonal degeneration are associated with memory, attentional, and executive functioning in early-stage parkinson’ s disease. International journal of molecular sciences 21: 8519

54 Oosterveld LP, Verberk IM, Majbour NK, El-Agnaf OM, Weinstein HC, Berendse HW, Teunissen CE, van de Berg WD (2020) CSF or serum neurofilament light added to α-synuclein panel discriminates Parkinson’s from controls. Movement Disorders 35: 288–295

55 Petzold A (2005) Neurofilament phosphoforms: surrogate markers for axonal injury, degeneration and loss. Journal of the neurological sciences 233: 183–198

56 Postuma RB, Berg D, Stern M, Poewe W, Olanow CW, Oertel W, Obeso J, Marek K, Litvan I, Lang AE (2015) MDS clinical diagnostic criteria for Parkinson’s disease. Movement disorders 30: 1591–1601

57 Sampedro F, Kulisevsky J (2022) Intracortical surface-based MR diffusivity to investigate neurologic and psychiatric disorders: a review. Journal of Neuroimaging 32: 28–35

58 Sampedro F, Martínez-Horta S, Marín-Lahoz J, Pagonabarraga J, Kulisevsky J (2019) Longitudinal intracortical diffusivity changes in de-novo Parkinson’s disease: a promising imaging biomarker. Parkinsonism & Related Disorders 68: 22–25

59 Sampedro F, Pérez-González R, Martínez-Horta S, Marín-Lahoz J, Pagonabarraga J, Kulisevsky J (2020) Serum neurofilament light chain levels reflect cortical neurodegeneration in de novo Parkinson’s disease. Parkinsonism & Related Disorders 74: 43–49

60 Steenwijk MD, Pouwels PJ, Daams M, van Dalen JW, Caan MW, Richard E, Barkhof F, Vrenken H (2013) Accurate white matter lesion segmentation by k nearest neighbor classification with tissue type priors (kNN-TTPs). NeuroImage: Clinical 3: 462–469

61 Taylor KI, Sambataro F, Boess F, Bertolino A, Dukart J (2018) Progressive decline in gray and white matter integrity in de novo Parkinson’s disease: an analysis of longitudinal Parkinson progression markers initiative diffusion tensor imaging data. Frontiers in aging neuroscience 10: 318

62 Thal DR, Ghebremedhin E, Rub U, Yamaguchi H, Del Tredici K, Braak H (2002) Two types of sporadic cerebral amyloid angiopathy. J Neuropathol Exp Neurol 61: 282–293 Doi 10.1093/jnen/61.3.282

63 Thal DR, Rüb U, Orantes M, Braak H (2002) Phases of Aβ-deposition in the human brain and its relevance for the development of AD. Neurology 58: 1791–1800

64 Tsuboi Y, Uchikado H, Dickson DW (2007) Neuropathology of Parkinson’s disease dementia and dementia with Lewy bodies with reference to striatal pathology. Parkinsonism & related disorders 13: S221–S224

65 Watson R, Blamire AM, Colloby SJ, Wood JS, Barber R, He J, O’Brien JT (2012) Characterizing dementia with Lewy bodies by means of diffusion tensor imaging. Neurology 79: 906–914

66 Weintraub D, Doshi J, Koka D, Davatzikos C, Siderowf AD, Duda JE, Wolk DA, Moberg PJ, Xie SX, Clark CM (2011) Neurodegeneration across stages of cognitive decline in Parkinson disease. Archives of neurology 68: 1562–1568

67 Yong AC, Tan YJ, Ng EY, Lu Z, Ng SY, Chia NS, Choi X, Heng D, Neo SX, Xu Z (2020) Association between plasma neurofilament light chain levels and cognition in early Parkinson’s disease: Biomarkers (non-neuroimaging)/Plasma/Serum/Urine biomarkers. Alzheimer’s & Dementia 16: e040206

