## Supplementary figures for "Cortical distribution of neurofilaments associates with pathological hallmarks and MRI measures of atrophy and diffusivity in Parkinson’s disease"

**Author affiliations:**

Correspondence to: Irene Frigerio

Journal name: Acta Neuropathologica

### Supplementary figures

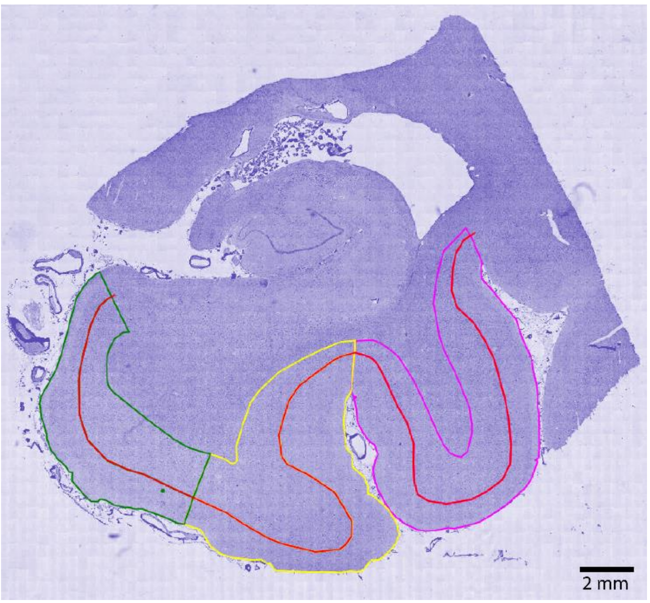


**Supplementary Fig. 1 Segmentation of entorhinal cortex, parahippocampal and fusiform gyrus in hippocampal section**. Hippocampal sections (here hematoxylin deconvoluted image) were segmented according to the method described by Adler et al. [1], where the entorhinal cortex (green), parahippocampal gyrus (yellow) and fusiform gyrus (pink) were delineated to match the MRI derived ROIs. Briefly, the entorhinal cortex was delineated from the end of the parasubiculum until layer IV started being visible [4]; the parahippocampal gyrus started from this point and ended at the collateral sulcus; the fusiform gyrus started at this point and ended at the inferior temporal sulcus. The line separating superficial (I-III) and deep (IV-VI) cortical layers can be seen in red in the figure. Note that the entorhinal cortex was subdivided into layers I-III (superficial) and lamina dissecans plus layers V-VI (deep) [35].

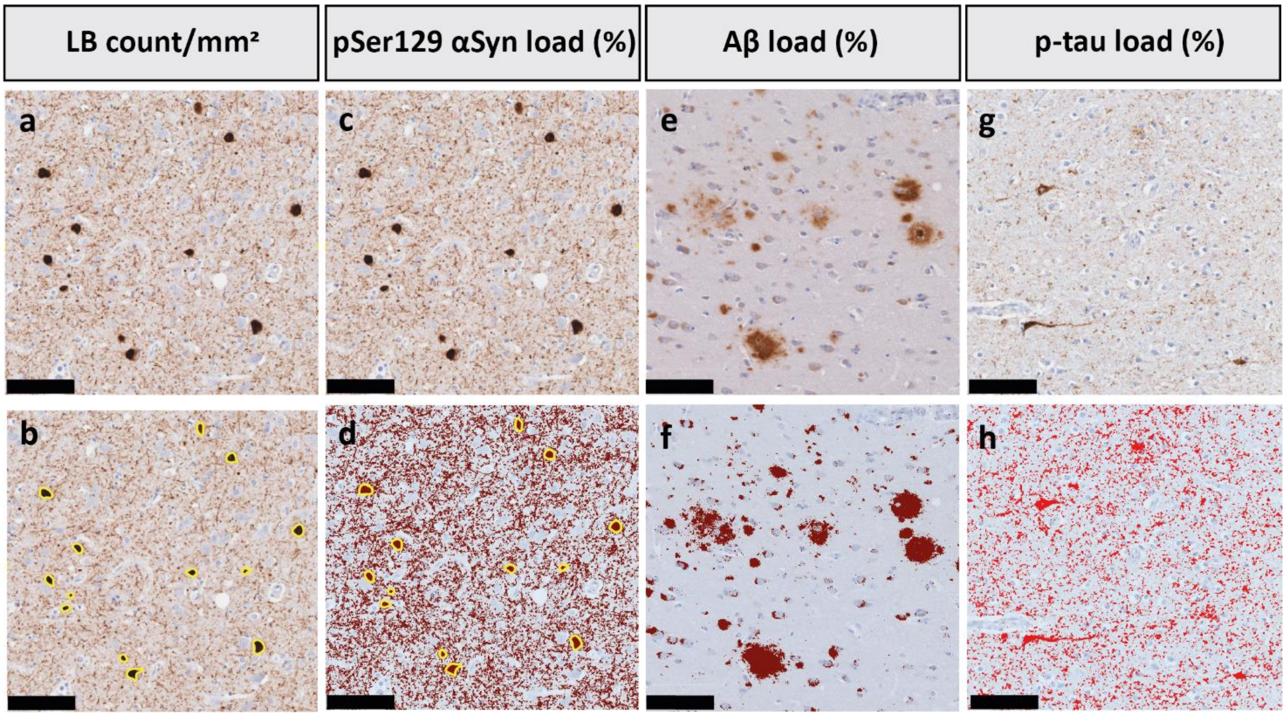


**Supplementary Fig. 2 Quantification of pathology load using QuPath [2] algorithms**. The DAB immunoreactive area of **(a,c)** pSer129-αSyn, **(e)** Aβ and **(g)** p-tau, and their (QuPath) quantification **(b, d, f, h**, respectively) are shown. In particular, **b)** shows the results of the object classifier detecting pSer129-αSyn-positive Lewy Bodies (LBs), encircled in yellow. **d)** shows the pixel classifier detecting pSer129-αSyn positive staining (in dark red), including all pSer129-αSyn positive morphologies except LBs (in yellow), such as Lewy neurites, and dot-like structures [5]. **f)** shows the result of the pixel classifier detecting Aβ positive staining (in dark red). **h)** shows the pixel classifier detecting p-tau positive staining (in red). The scale bar in each image corresponds to 100 µm. **Legend**: *Aβ: amyloid beta; LB: Lewy Body; pSer129-αSyn: phosphorylated Ser129 alpha synuclein; p-tau: phosphorylated tau.*

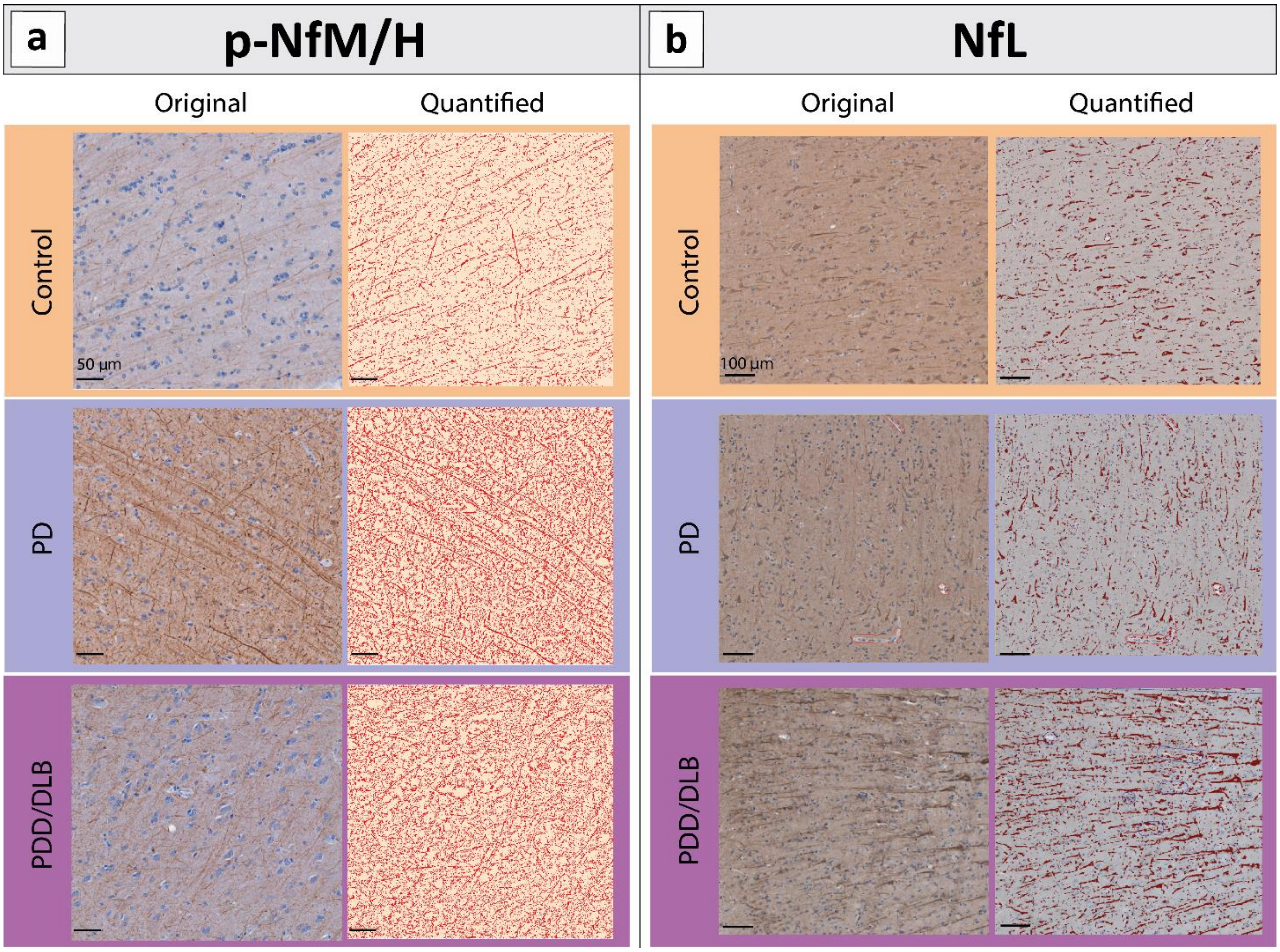


**Supplementary Fig. 3 Photographs of representative sections of controls, PD and PDD/DLB donors stained for neurofilament NfL and p-NfM/H**. Image before (original) and after processing with QuPath [2] *in-house* pixel classifier are shown. Representative images (parahippocampal gyrus) of **a** p-NfM/H and **b** NfL immunoreactivity are shown for controls, PD and PDD/DLB. Morphologically, p-NfM/H staining was observed in axons, while NfL staining was seen in the neuronal somas and its processes. P-NfM/H immunoreactivity was increased in both PD and PDD/DLB compared to controls, while NfL immunoreactivity was increased in only PDD/DLB compared to controls. **Legend**: *DLB: Dementia with Lewy Bodies; NfL: neurofilament light chain; PD: Parkinson's disease; PDD: Parkinson's disease dementia; p-NfM/H: phosphorylated neurofilament medium and heavy chain.*

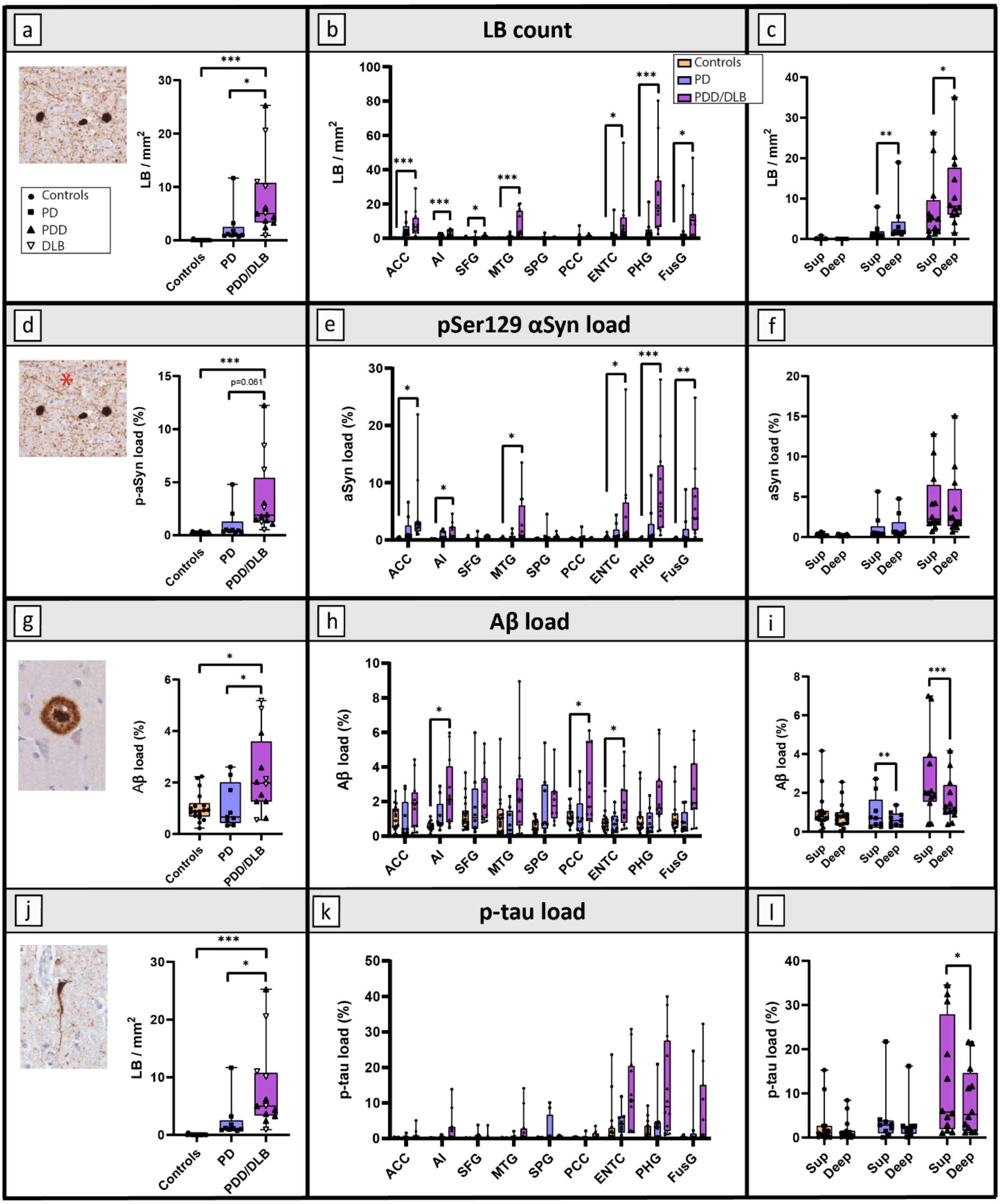


**Supplementary Fig. 4 Cortical pathology load in controls, PD and PDD/DLB donors**. **a, b** and **c** the pSer129-αSyn-positive LB count (LB/mm$^2$), **d, e** and **f** pSer129-αSyn load (indicated by red asterisk in figure; %area), **g, h** and **i** Aβ load (%area), **j, k** and **l** and p-tau load (%area) are shown for controls, PD and PDD/DLB groups. The *left column* shows the overall pathology load across all the cortical regions examined; every data point represents a donor averaged across all cortical regions, and the clinical groups are indicated by shapes in **a**. The *middle column* shows the regional pathology load in all the 9 cortical regions across groups. The *right column* shows the pathology load in superficial (sup; layer I-III) and deep cortical layers (layer IV-VI) in each group for each outcome measure. **c** and **f** show the cortical layer load across all the regions examined except for the superior parietal gyrus and the posterior cingulate gyrus (therefore including only the regions that showed significant changes across groups), while **i** and **l** show Aβ and p-tau load in superficial (layer I-III) and deep cortical layers (layer IV-VI) of the cortical regions near the hippocampus (entorhinal cortex, parahippocampal gyrus and fusiform gyrus), respectively, as these are among the first regions to be affected by Alzheimer's disease pathology [3, 6]. **Legend**: **p<0.05, **p<0.01, ***p<0.001 when compared to controls; Aβ: amyloid beta; ACC: anterior cingulate cortex; AI: anterior insula; DLB: Dementia with Lewy Bodies; ENTC: entorhinal cortex; FusG: fusiform gyrus; LB: Lewy Body; MTG: middle temporal gyrus; PCC: posterior cingulate gyrus; PHG: parahippocampal gyrus; PD: Parkinson's disease; PDD: Parkinson's disease dementia; pSer129-αSyn: phosphorylated Ser129 alpha synuclein; p-tau: phosphorylated tau; SFG: superior frontal gyrus; SPG: superior parietal gyrus; sup: superficial.*

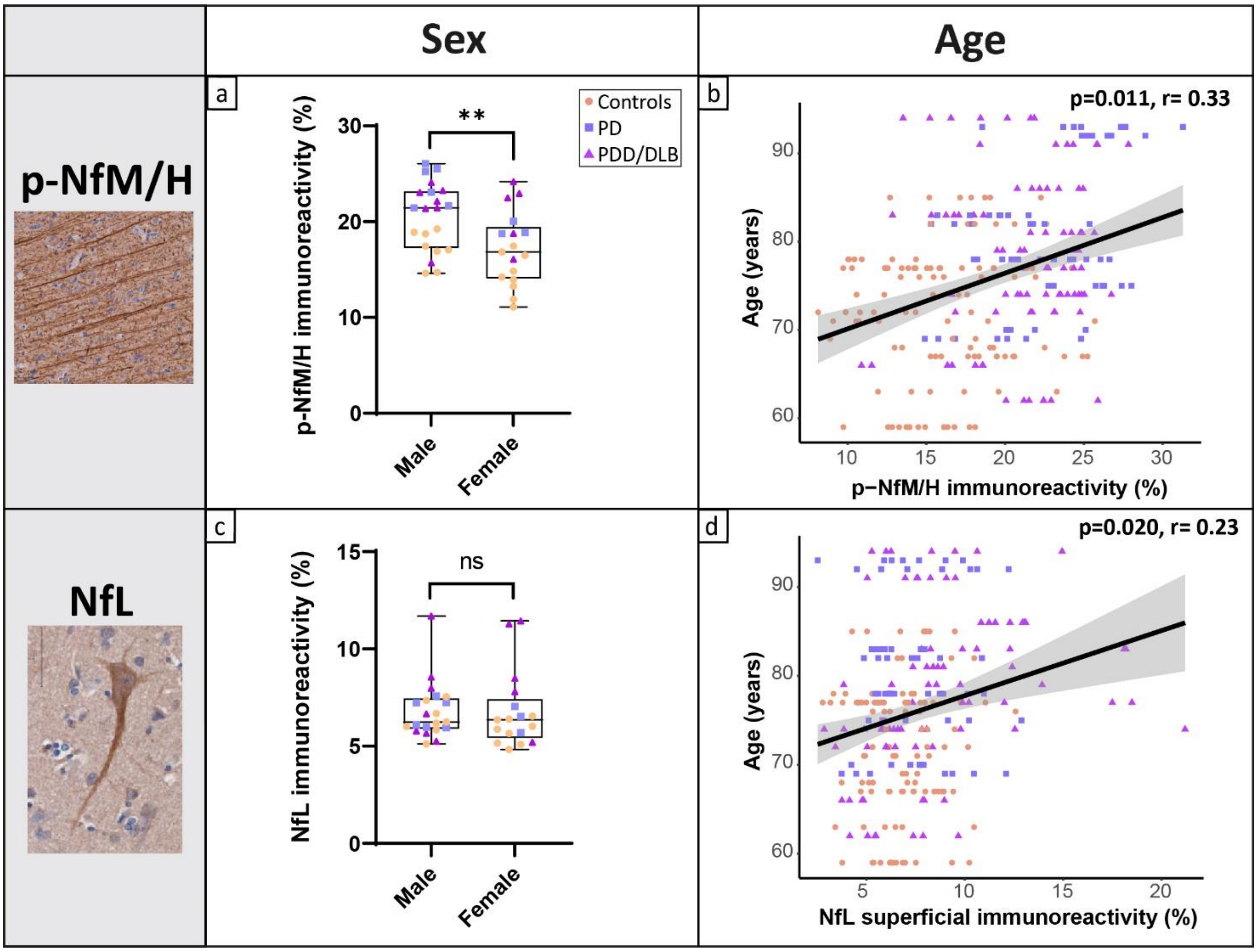


**Supplementary Fig. 5 Differences in cortical neurofilaments immunoreactivity in sex, and neurofilaments association with age.** The *left column* shows **a** p-NfM/H and **c** NfL immunoreactivity in male versus female donors across the whole cohort. The *right column* shows the correlation between age at death and **b** p-NfM/H and **d** NfL immunoreactivity**.** P-NfM/H was significantly higher in males than females in the entire cohort, while there was no difference in NfL immunoreactivity in gender. Both neurofilaments immunoreactivity levels increased with age at death (regarding NfL, specifically NfL immunoreactivity in superficial cortical layers). In **a** and **c** every data point represents a donor, while in **b** and **d** every data point represents a brain area of a donor, and the regression line with the standard error are shown. Groups are indicated by shapes in **a**. **Legend**: ***p<0.01 when compared to controls; DLB: Dementia with Lewy Bodies; NfL: neurofilament light chain; ns: not significant; PD: Parkinson's disease; PDD: Parkinson's disease dementia; p-NfM/H: phosphorylated neurofilament medium and heavy chain.*

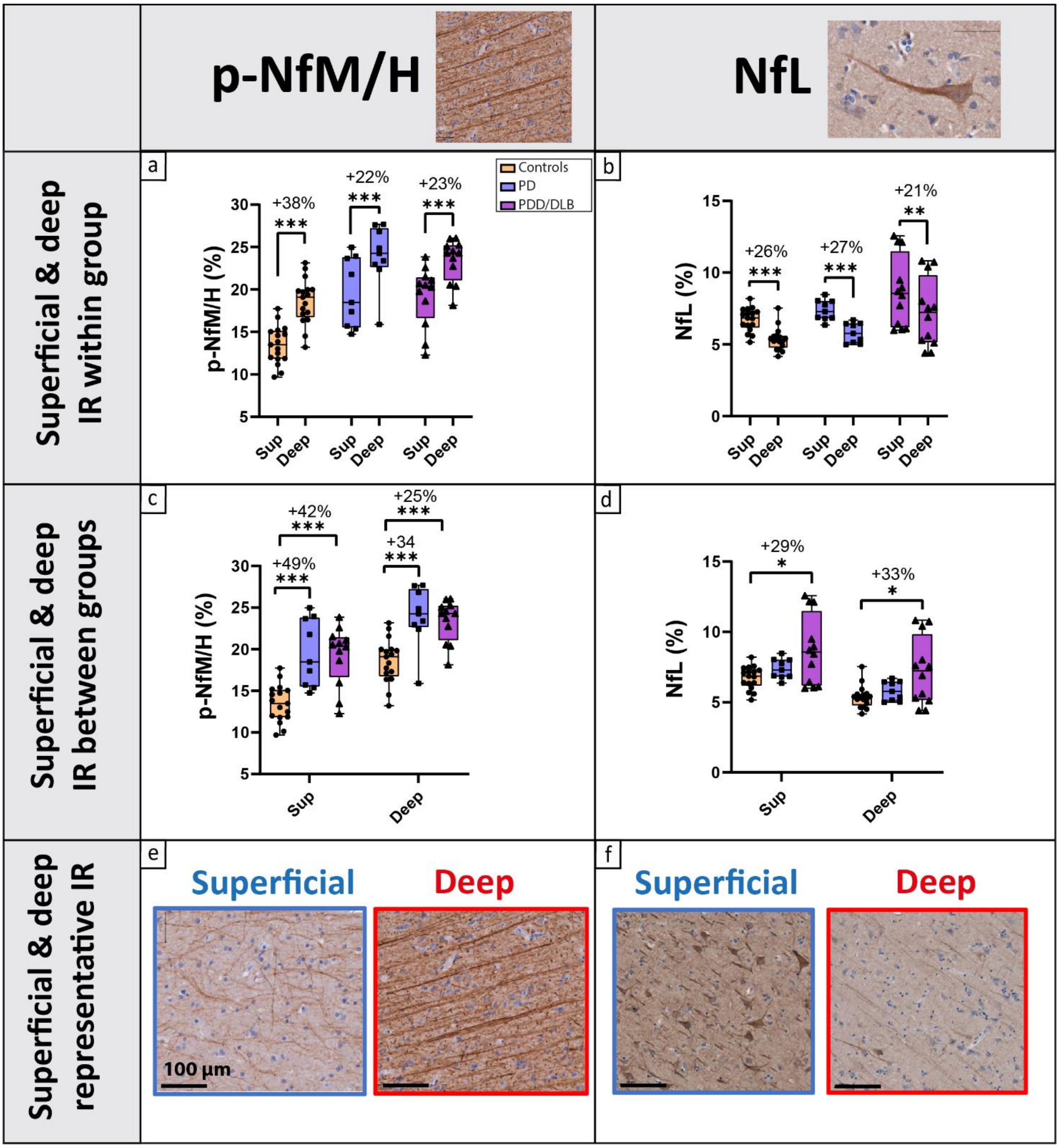


**Supplementary Fig. 6 Cortical neurofilaments immunoreactivity (IR) in superficial (layer I-III) and deep cortical layers (layer IV-VI).** The *left column* shows p-NfM/H and *right column* shows NfL immunoreactivity behaviour in superficial (layers I-III) and deep (layers IV-VI) cortical layers. Specifically,

**a** p-NfM/H and **b** NfL immunoreactivity in superficial (layer I-III) and deep cortical layers (layer IV-VI) *within* group (percentage differences shown). **c** p-NfM/H and **d** NfL load in superficial and deep cortical layers *between* groups (percentage differences when compared to controls shown). Each data point represents a donor. Groups are indicated by colors in **a**. **e** Representative images of p-NfM/H and **F)** NfL immunoreactivity in superficial and deep cortical layers. Overall, P-NfM/H immunoreactivity was higher in deep compared to superficial cortical layers in all groups, while NfL immunoreactivity showed the opposite trend, being more abundant in superficial than in deep cortical layers in all groups. **Legend**: **p<0.05, **p<0.01, ***p<0.001 when compared to controls; DLB: Dementia with Lewy Bodies; IR: immunoreactivity; NfL: neurofilament light chain; PD: Parkinson's disease; PDD: Parkinson's disease dementia; p-NfM/H: phosphorylated neurofilament medium and heavy chain; sup: superficial.*

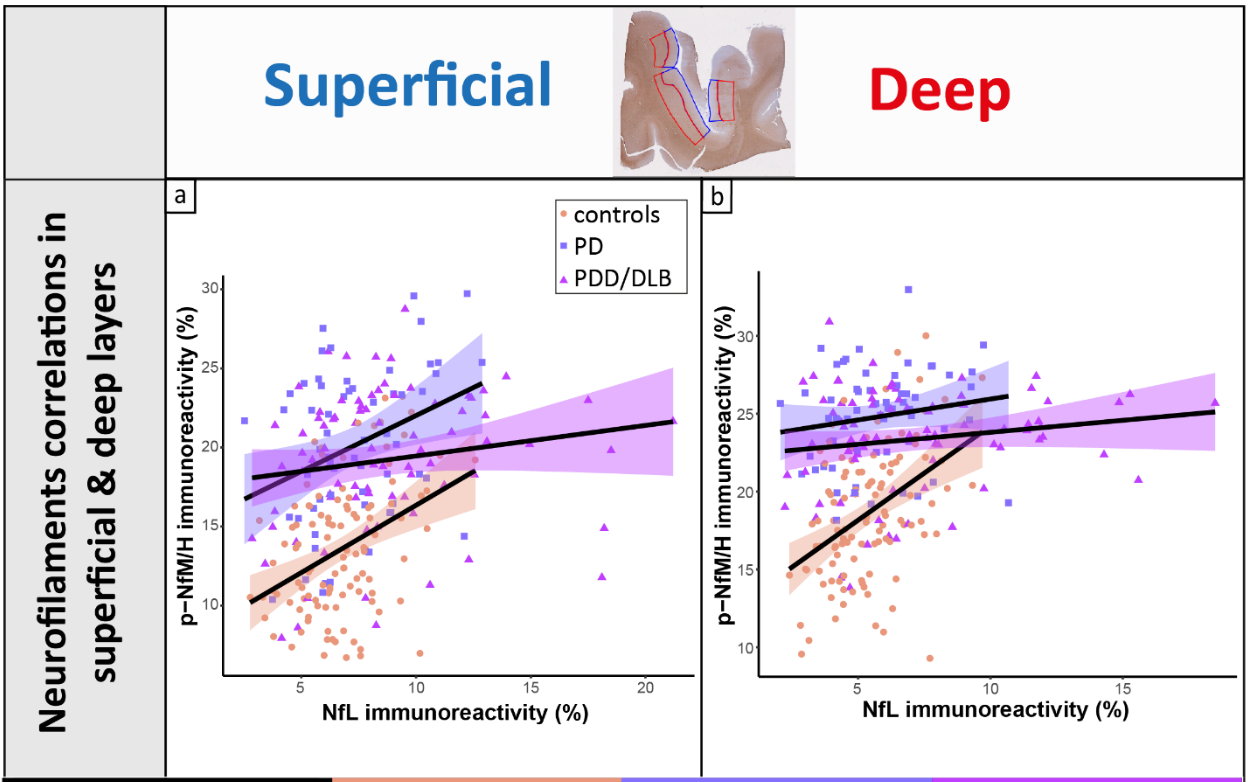


| | Controls | PD | PDD/DLB |
|---|---|---|---|
| **NfL & p-NfM/H** | | | |
| **Entire CORTEX** | **p<0.001, r = 0.39** | p=0.054, r = 0.19 | p=0.089, r = 0.17 |
| ***Superficial layers*** | **p<0.001, r = 0.38** | **p=0.006, r = 0.25** | **p<0.001, r = 0.30** |
| ***Deep layers*** | **p<0.001, r = 0.34** | p=0.242, r = 0.13 | p=0.061, r = 0.20 |

**Supplementary Fig. 7 Cortical neurofilaments correlation in superficial (layer I-III) and deep cortical layers (layer IV-VI).** Correlations between p-NfM/H and NfL immunoreactivity in **a** superficial (layer I-III) and **b** deep cortical layers (layer IV-VI), indicated also in the section in the top row (blue annotations correspond to superficial cortical layers, and red annotations to deep cortical layers). Each data point represents a brain area of a donor. Groups are indicated by colors and shapes in **a**. Underneath, the table with the p-NfM/H and NfL correlations in the entire cortex (layer I-VI), superficial (layer I-III) and deep cortical layers (layer IV-VI) within groups. For each correlation, p-value and correlation coefficient are shown.

When the correlation is significant, it is highlighted in bold, and the strength of the correlation is given by a colour (dark green moderate correlation, light green is weak correlation). Correlations coloured in light blue are trends. Overall, p-NfM/H and NfL immunoreactivity moderately correlated in controls independently of the layer, but this correlation was lost in PD and PDD/DLB. Specifically, the correlation was lost in the deep cortical layers of both PD and PDD/DLB donors, while it was still present in the superficial layers of both PD and PDD/DLB. **Legend**: *DLB: Dementia with Lewy Bodies; NfL: neurofilament light chain; PD: Parkinson's disease; PDD: Parkinson's disease dementia; p-NfM/H: phosphorylated neurofilament medium and heavy chain.*

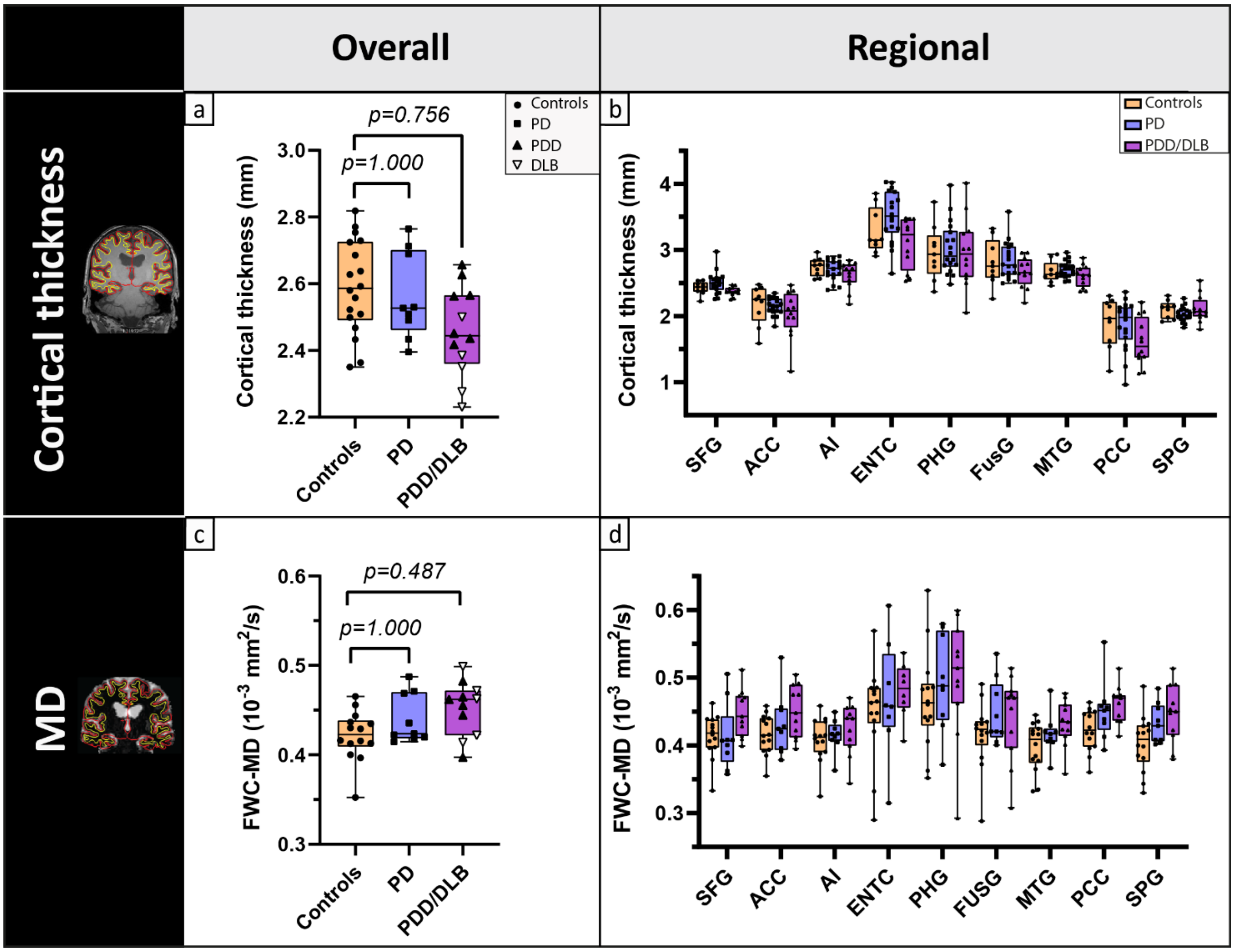


**Supplementary Fig. 8 Cortical thickness and MD do not differ between controls, PD and PDD/DLB donors**. **a** and **b** differences in cortical thickness and **c** and **d** cortical MD across the regions of interest selected for this study are shown. Specifically, **a** and **c** show the MRI outcome measure across all regions, where every data point represents a donor (clinical groups shapes indicated in **a**, and **b** and **d** show the regional changes in the outcome measure (groups colors indicated in **b**). No difference were found in both cortical thickness and cortical MD between groups. **Legend**: *ACC: anterior cingulate cortex; AI: anterior insula; DLB: Dementia with Lewy Bodies; ENTC: entorhinal cortex; FUSG: fusiform gyrus; FWC-MD: free*

*water corrected mean diffusivity; MD: mean diffusivity; MTG: middle temporal gyrus; PCC: posterior cingulate gyrus; PHG: parahippocampal gyrus; PD: Parkinson's disease; PDD: Parkinson's disease dementia; SFG: superior frontal gyrus; SPG: superior parietal gyrus.*

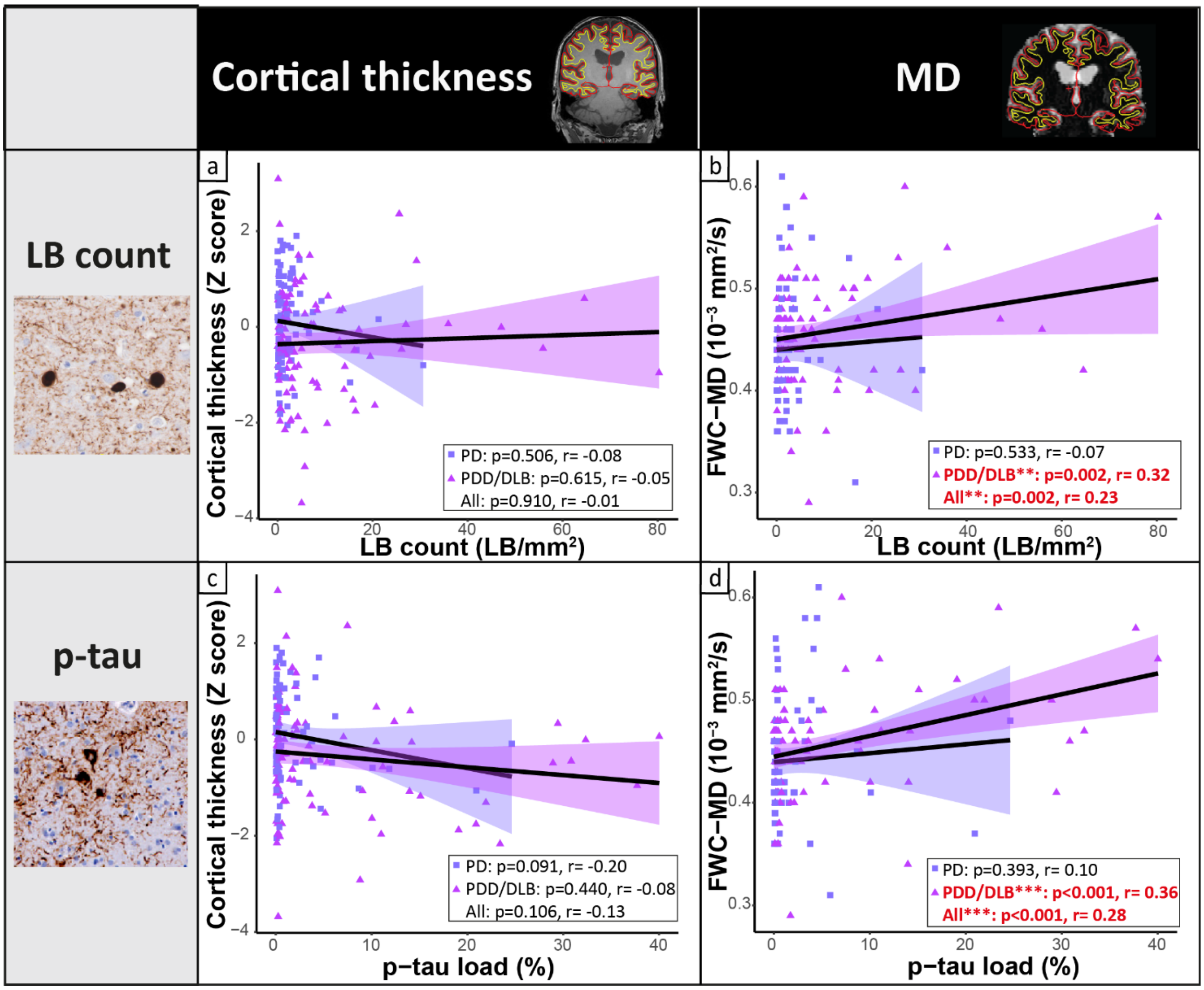


**Supplementary Fig. 9 Cortical MD, but not cortical thickness, correlates with pSer129-αSyn and p-tau pathology**. Correlations between **a** and **b** pSer129-αSyn-positive LB count and **c** and **d** p-tau load with MRI derived-cortical thickness on the left, and DTI-derived cortical MD on the right**.** Higher LB count and p-tau load associated with increased cortical MD, and not with cortical thickness. Every data point represents a brain area of a donor, and the groups are indicated by shapes in the figure. For each group, the regression line is shown, together with its colour-coded standard error (purple for PD and pink for PDD/DLB), and the p-values and correlation coefficient (r) of the correlation (for PD, PDD/DLB, and all data points together). Significant correlations are highlighted in red bold in the box in every panel. **Legend**: correlation significant at **p<0.05, **p<0.01, ***p<0.001; DLB: Dementia with Lewy Bodies; FWC-MD: free-water corrected mean diffusivity; LB: Lewy Body; MD: mean diffusivity; PD: Parkinson's disease; PDD: Parkinson's disease dementia; p-tau: phosphorylated tau.*

### References supplementary figures


1 Adler DH, Pluta J, Kadivar S, Craige C, Gee JC, Avants BB, Yushkevich PA (2014) Histology-derived volumetric annotation of the human hippocampal subfields in postmortem MRI. Neuroimage 84: 505-523

2 Bankhead P, Loughrey MB, Fernandez JA, Dombrowski Y, McArt DG, Dunne PD, McQuaid S, Gray RT, Murray LJ, Coleman HGet al (2017) QuPath: Open source software for digital pathology image analysis. Sci Rep 7: 16878 Doi 10.1038/s41598-017-17204-5

3 Braak H, Alafuzoff I, Arzberger T, Kretzschmar H, Del Tredici K (2006) Staging of Alzheimer disease-associated neurofibrillary pathology using paraffin sections and immunocytochemistry. Acta neuropathologica 112: 389-404

4 Insausti R, Munoz-Lopez M, Insausti AM, Artacho-Perula E (2017) The Human Periallocortex: Layer Pattern in Presubiculum, Parasubiculum and Entorhinal Cortex. A Review. Front Neuroanat 11: 84 Doi 10.3389/fnana.2017.00084

5 Kovacs GG, Wagner U, Dumont B, Pikkarainen M, Osman AA, Streichenberger N, Leisser I, Verchère J, Baron T, Alafuzoff I (2012) An antibody with high reactivity for disease-associated α-synuclein reveals extensive brain pathology. Acta neuropathologica 124: 37-50

6 Thal DR, Rüb U, Orantes M, Braak H (2002) Phases of Aβ-deposition in the human brain and its relevance for the development of AD. Neurology 58: 1791-1800
