## Supplementary material for "Cortical distribution of neurofilaments associates with pathological hallmarks and MRI measures of atrophy and diffusivity in Parkinson’s disease"

#### Pathological pSer129-αSyn quantification

To determine the load of pSer129-αSyn pathology, a script was developed in QuPath [2] that first identified and counted LB-like structures (both intra- and extracellular) and then determined the %area load of the non-LB pSer129-αSyn pathology. In this study, the LB-like structures count is referred to as LB count, but it is important to mention that the LB count might also occasionally include corpora amylaceas [1, 5] since they are similar in morphology to LBs and positive to pSer129-αSyn staining (even though they are located mostly in superficial rather than deep cortical layers). That is why, in our cohort, control cases showed a few LB-like structures, which corresponded not to LBs but to corpora amylaceas (checked by I.F.). To detect LBs, an object classifier was created based on the DAB signal, and the object diameter was defined at 5-25 µm (or an area between 28.27 and 490.9 $\mu m^2$), based on the definition of LBs [3, 4]. After counting LB-like structures, the LB detections were excluded from the ROIs and pixel classification was used to quantify the %area load of non-LB pSer129-αSyn pathology, which includes morphologies such Lewy Neurites and a variety of morphological inclusions [3] (**Supplementary Figure 1).**

### References supplementary material


1 Augé E, Cabezón I, Pelegrí C, Vilaplana J (2017) New perspectives on corpora amylacea in the human brain. Scientific Reports 7: 1-10

2 Bankhead P, Loughrey MB, Fernandez JA, Dombrowski Y, McArt DG, Dunne PD, McQuaid S, Gray RT, Murray LJ, Coleman HGet al (2017) QuPath: Open source software for digital pathology image analysis. Sci Rep 7: 16878 Doi 10.1038/s41598-017-17204-5

3 Kovacs GG, Wagner U, Dumont B, Pikkarainen M, Osman AA, Streichenberger N, Leisser I, Verchère J, Baron T, Alafuzoff I (2012) An antibody with high reactivity for disease-associated α-synuclein reveals extensive brain pathology. Acta neuropathologica 124: 37-50

4 Moors TE, Maat CA, Niedieker D, Mona D, Petersen D, Timmermans-Huisman E, Kole J, El-Mashtoly SF, Spycher L, Zago W (2021) The subcellular arrangement of alpha-synuclein proteoforms in the Parkinson’s disease brain as revealed by multicolor STED microscopy. Acta neuropathologica 142: 423-448

5 Navarro PP, Genoud C, Castaño-Díez D, Graff-Meyer A, Lewis AJ, de Gier Y, Lauer ME, Britschgi M, Bohrmann B, Frank S (2018) Cerebral Corpora amylacea are dense membranous labyrinths containing structurally preserved cell organelles. Scientific reports 8: 1-13
