## Supplementary tables for "Cortical distribution of neurofilaments associates with pathological hallmarks and MRI measures of atrophy and diffusivity in Parkinson’s disease"

#### Supplementary Table 1. Donor characteristics.

| Case number | Clinical diagnosis | Sex | Age at death (years) | Disease duration (years) | PMD (hr:min) | Cause of death | 3DT1 scan | DTI scan | IHC | NBV (L) | NGMV (L) | ABC score [5] | Thal phase [6] | Braak NFT stage [1] | Braak LB stage [2] |
|---|---|---|---|---|---|---|---|---|---|---|---|---|---|---|---|
| **Controls** | | | | | | | | | | | | | | | |
| 1 | Ctrl | M | 68 | - | 8:30 | Euthanasia | x | | x | 1,62 | 0,86 | A1B1C0 | 2 | 1 | 0 |
| 2 | Ctrl | F | 63 | - | 8:10 | Euthanasia | x | x | x | 1,53 | 0,75 | A0B0C0 | 0 | 0 | 0 |
| 3 | Ctrl | M | 82 | - | 10:40 | Liver cirrhosis | x | x | x | 1,40 | 0,68 | A1B1C0 | 1 | 1 | 0 |
| 4 | Ctrl | M | 85 | - | 9:22 | Euthanasia | x | x | x | 1,36 | 0,68 | A1B1C0 | 1 | 1 | 0 |
| 5 | Ctrl | M | 67 | - | 7:35 | Euthanasia | x | x | x | 1,51 | 0,74 | A1B1C0 | 1 | 2 | 0 |
| 6 | Ctrl | F | 76 | - | 7:50 | Euthanasia | x | x | x | 1,47 | 0,78 | A1B1C0 | 2 | 1 | 0 |
| 7 | Ctrl | M | 67 | - | 8:10 | Liver cirrhosis | x | x | x | 1,45 | 0,74 | A1B1C0 | 1 | 1 | 0 |
| 8 | Ctrl | F | 77 | - | 9:00 | Euthanasia | x | x | x | 1,41 | 0,74 | A1B1C0 | 1 | 2 | 0 |
| 9 | Ctrl | F | 87 | - | 8:35 | Urinary tract infection | x | | x | 1,37 | 0,65 | A0B1C0 | 0 | 1 | 0 |
| 10 | Ctrl | F | 72 | - | 7:20 | Heart failure | x | | x | 1,52 | 0,80 | A0B0C0 | 0 | 0 | 0 |
| 11 | Ctrl | F | 69 | - | 12:45 | Pulmonary embolism | x | | x | 1,39 | 0,71 | A1B1C0 | 1 | 1 | 1 |
| 12 | Ctrl | M | 59 | - | 8:00 | Euthanasia | x | | x | 1,49 | 0,76 | A1B1C0 | 2 | 1 | 0 |
| 13 | Ctrl | M | 77 | - | 11:25 | Pneumonia | x | x | x | 1,49 | 0,72 | A1B1C0 | 1 | 1 | 0 |
| 14 | Ctrl | F | 78 | - | 10:00 | Unknown | x | x | x | 1,51 | 0,77 | A1B1C0 | 1 | 1 | 1 |
| 15 | Ctrl | F | 77 | - | 4:35 | Unknown | x | x | x | 1,46 | 0,76 | A1B1C0 | 2 | 2 | 2 |
| 16 | Ctrl | F | 59 | - | 8:10 | Euthanasia | x | x | x | 1,46 | 0,78 | A0B0C0 | 0 | 0 | 0 |
| 17 | Ctrl | F | 71 | - | 6:50 | Lung carcinoma | x | x | x | 1,47 | 0,76 | A1B1C0 | 2 | 1 | 0 |
| 18 | Ctrl | M | 74 | - | 10:20 | Euthanasia | x | x | x | 1,39 | 0,70 | A2B1C0 | 3 | 2 | 0 |
| **PD** | | | | | | | | | | | | | | | |
| 1 | PD | F | 83 | 22 | 10:35 | Euthanasia | x | x | x | 1,42 | 0,68 | A0B1C0 | 0 | 1 | 5 |

| | | | | | | | | | | | | | | | |
|---|---|---|---|---|---|---|---|---|---|---|---|---|---|---|---|
| 2 | PD | F | 69 | 15 | 07:05 | Aspiration pneumonia | x | x | x | 1,51 | 0,73 | A1B1C0 | 2 | 2 | 6 |
| 3 | PD | F | 82 | 17 | 09:17 | Aspiration pneumonia | x | x | x | 1,44 | 0,73 | A1B1C0 | 2 | 2 | 6 |
| 4 | PD | M | 78 | 17 | 07:15 | Euthanasia | x | x | x | 1,45 | 0,76 | A1B1C0 | 1 | 2 | 6 |
| 5 | PD | M | 92 | 16 | 10:10 | Myocardial infarction | x | x | x | 1,38 | 0,70 | A2B2C1 | 3 | 3 | 4 |
| 6 | PD | M | 75 | 20 | 4:55 | End-stage PD | x | x | x | 1,41 | 0,74 | A2B1C0 | 3 | 2 | 6 |
| 7 | PD | M | 78 | 20 | 3:30 | End-stage PD | x | x | x | 1,25 | 0,63 | A1B1C0 | 1 | 2 | 6 |
| 8 | PD | M | 70 | 8 | 6:55 | Euthanasia | x | x | x | 1,53 | 0,78 | A1B1C0 | 1 | 1 | 6 |
| 9 | PD | M | 93 | 23 | 10:40 | Euthanasia | x | x | x | 1,50 | 0,72 | A1B2C0 | 2 | 4 | 6 |
| **PDD/ DLB** | | | | | | | | | | | | | | | |
| 1 | PDD | F | 83 | 16 | 10:40 | End-stage PDD | x | x | x | 1,33 | 0,65 | A3B2C2 | 4 | 4 | 6 |
| 2 | DLB | M | 66 | 4 | 08:00 | Euthanasia | x | x | x | 1,49 | 0,73 | A3B2C2 | 4 | 4 | 6 |
| 3 | PDD | F | 94 | 10 | 06:50 | Femur fracture | x | x | x | 1,33 | 0,68 | A3B2C2 | 4 | 4 | 6 |
| 4 | PDD | F | 74 | 12 | 08:10 | Euthanasia | x | x | x | 1,33 | 0,71 | A2B1C0 | 3 | 2 | 6 |
| 5 | DLB | M | 72 | 5 | 07:10 | Euthanasia | x | x | x | 1,42 | 0,72 | A2B1C2 | 3 | 2 | 6 |
| 6 | DLB | M | 77 | 7 | 07:15 | End-stage DLB | x | x | x | 1,45 | 0,75 | A1B2C0 | 1 | 3 | 6 |
| 7 | PDD | M | 79 | 21 | 09:25 | Subarachnoid bleeding | x | x | x | 1,38 | 0,71 | A2B1C1 | 3 | 2 | 6 |
| 8 | PDD | M | 62 | 18 | 05:10 | End-stage PDD | x | | x | 1,37 | 0,71 | A1B1C0 | 1 | 2 | 6 |
| 9 | PDD | M | 74 | 8 | 8:50 | Aspiration pneumonia | x | x | x | 1,29 | 0,62 | A1B2C0 | 2 | 3 | 6 |
| 10 | PDD | F | 81 | 19 | 5:30 | End-stage PDD | x | x | x | 1,35 | 0,66 | A2B1C0 | 3 | 2 | 6 |
| 11 | DLB | M | 91 | 3 | 4:30 | Euthanasia | x | x | x | 1,48 | 0,72 | A2B2C2 | 3 | 4 | 6 |
| 12 | DLB | F | 86 | 6 | 7:00 | Dehydration, cachexia | x | x | x | 1,31 | 0,69 | A3B2C2 | 5 | 4 | 6 |

**Legend**: *3DT1: post-mortem in-situ MRI 3DT1 weighted scan; Ctrl: control; DLB: Dementia with Lewy Bodies; DTI: diffusion tensor imaging; F: female; hr: hour; IHC: immunohistochemistry; L: liter; LB: Lewy Body; M: male; min: minute; NBV: normalized brain volume; NFT: neurofibrillary tangles; NGMV: normalized grey matter volume; PD: Parkinson's disease; PDD: Parkinson's disease dementia; PMD: post-mortem delay.*

**Supplementary Table 2. Concatenated labels from Lausanne atlas for the regions of interest of this study.**

| Region of interest | Labels Lausanne atlas [3, 4] |
|---|---|
| Superior frontal | Superiorfrontal_4 + Superiorfrontal_5 + Superiorfrontal_6 |
| Anterior cingulate | Caudalanteriorcingulate_2 + Caudalanteriorcingulate_3 |
| Posterior cingulate | Posteriorcingulate_3 + Posteriorcingulate_4 |
| Superior parietal | Superiorparietal_1 + Superiorparietal_2 + Superiorparietal_3 |
| Parahippocampal | Parahippocampal_3 |
| Entorhinal | Entorhinal_1 |
| Fusiform | Fusiform_7 |
| Middle temporal | Middletemporal_5 + Middletemporal_7 |
| Anterior insula | Insula_3 + Insula_4 |

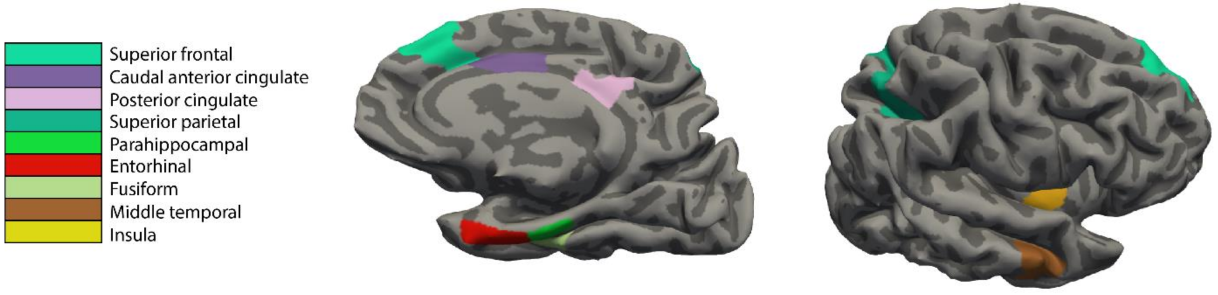

**Supplementary Table 3. Information on primary and secondary antibodies.**

| *Antibody* | **Antigen** | **Species** | **Origin details** | **Dilution** | **Incubation time** | **Antigen retrieval** | **Detection method** |
|---|---|---|---|---|---|---|---|
| ***Primary antibodies*** | | | | | | | |
| *pSer129 aSyn, clone EP1536Y* | Alpha synuclein phosphorylated at Ser129 | Rabbit igG | Abcam, Cambridge, UK | 1:8000 for BF<br>1:4000 for FL | 4°C o.n. | Tris EDTA buffer (pH 9.0) in steam cooker | EnVision (HRP) |
| *Aβ, clone 4G8* | Aβ amino acid sequence 17-24 | Mouse igG2b | BioLegend, San Diego, USA | 1:8000 | 4°C o.n. | Citrate buffer (pH 6.0) in steam cooker | EnVision (HRP) |
| *p-tau, clone AT8* | Tau phosphorylated at Ser202 and Thr205 | Mouse igG1 | ThermoFisher, Pittsburgh, USA | 1:800 for BF<br>1:800 for FL | 4°C o.n. | Citrate buffer (pH 6.0) in steam cooker | EnVision (HRP) |
| *NfL* | Immunogen corresponds to AA 1 to 284 (with AA 200-292 missing) | Rabbit igG | Synaptic systems, Göttingen, Germany | 1:600 for BF<br>1:200 for FL | 4°C o.n. | Tris EDTA buffer (pH 9.0) in steam cooker | EnVision (HRP) |
| *SMI312* | Mixture of monoclonal antibodies directed against phosphorylated NfM and NfH. | Mouse igG1 | BioLegend, San Diego, USA | 1:500 for BF<br>1:500 for FL | 4°C o.n. | Tris EDTA buffer (pH 9.0) in steam cooker | EnVision (HRP) |
| ***Secondary antibodies*** | **Host species** | **Target species** | **Origin details** | **Dilution** | **Incubation time** | **Antigen retrieval** | **Detection method** |
| *DoAR Alexa 488* | Donkey IgG | Mouse | ThermoFisher, Pittsburgh, USA | 1:200 | 2 hrs at RT | / | fluorochrome |
| *DoAR Alexa 594* | Donkey IgG | Rabbit | ThermoFisher, Pittsburgh, USA | 1:200 | 2 hrs at RT | / | fluorochrome |

**Legend**: *BF: bright field; FL: fluorescence; hrs: hours; o.n.: overnight; RT: room temperature.*

**Supplementary Table 4. Correlations between pathological hallmarks.**

| | Controls | PD + PDD/DLB | PD | PDD/DLB |
|---|---|---|---|---|
| ***LB count (LB/mm$^2$)*** | | | | |
| *p-αSyn load (%)* | - | **p<0.001, r = 0.89** | **p<0.001, r = 0.84** | **p<0.001, r = 0.88** |
| *p-tau load (%)* | - | **p<0.001, r = 0.63** | p=0.611, r = 0.06 | **p<0.001, r = 0.68** |
| *Aβ load (%)* | | p=0.909, r = -0.01 | p=0.282, r = -0.13 | p=0.999, r = 0.00 |
| ***pSer129-αSyn load (%)*** | | | | |
| *p-tau load (%)* | - | **p<0.001, r = 0.57** | p=0.607, r = 0.07 | **p<0.001, r = 0.60** |
| *Aβ load (%)* | - | p=0.365, r = -0.05 | p=0.670, r = -0.05 | p=0.369, r = -0.06 |
| ***p-tau load (%)*** | | | | |
| *Aβ load (%)* | p=0.280, r = -0.01 | p=0.687, r = 0.02 | p=0.623, r = 0.05 | p=0.906, r = 0.01 |

For each correlation, p-value and correlation coefficients are shown. When the correlation is significant it is highlighted in bold. **Legend**: *Aβ: amyloid beta; DLB: Dementia with Lewy Bodies; LB: Lewy Body; PD: Parkinson's disease; PDD: Parkinson's disease dementia; pSer129-αSyn: phosphorylated Ser129 alpha synuclein; p-tau: phosphorylated tau.*

**Supplementary Table 5. Correlations of p-NfM/H immunoreactivity with pathology load in superficial (layer I-III) and deep cortical layers (layer IV-VI).**

| | **Controls** | **PD + PDD/DLB** | **PD** | **PDD/DLB** |
|---|---|---|---|---|
| ***p-NfM/H immunoreactivity (%)*** | | | | |
| *LB count (LB/mm$^2$) :* entire cortex | p=0.187, r = 0.15 | p=0.525, r = 0.07 | p=0.319, r = 0.15 | p=0.920, r = 0.01 |
| superficial | p=0.650, r = 0.04 | p=0.416, r = 0.08 | p=0.711, r = 0.06 | p=0.444, r = 0.11 |
| deep | p=0.214, r = -0.13 | p=0.567, r = 0.05 | p=0.132, r = 0.19 | p=0.907, r = -0.01 |
| *pSer129-αSyn load (%):* entire cortex | p=0.760, r = 0.03 | p=0.807, r = 0.03 | p=0.381, r = 0.13 | p=0.825, r = -0.03 |
| superficial | p=0.904, r = 0.01 | p=0.834, r = 0.02 | p=0.823, r = 0.03 | p=0.856, r = -0.02 |
| deep | p=0.370, r = 0.09 | p=0.886, r = 0.01 | *p=0.088, r = 0.22* | p=0.728, r = -0.04 |
| *p-tau load (%):* entire cortex | p=0.149, r = -0.18 | **p=0.018, r = 0.26** | *p=0.066, r = 0.32* | **p=0.043, r = 0.29** |
| superficial | p=0.707, r = -0.06 | p=0.704, r = 0.05 | p=0.168, r = 0.34 | p=0.608, r = -0.09 |
| deep | p=0.581, r = -0.11 | p=0.361, r = -0.13 | p=0.758, r = 0.06 | p=0.283, r = -0.22 |
| *Aβ load (%):* entire cortex | p=0.737, r = -0.03 | p=0.441, r = -0.06 | p=0.725, r = 0.05 | p=0.248, r = -0.12 |
| superficial | p=0.603, r = -0.06 | p=0.196, r = -0.10 | p=0.776, r = -0.04 | p=0.096, r = -0.18 |
| deep | p=0.892, r = 0.02 | p=0.784, r = -0.02 | p=0.683, r = -0.07 | p=0.964, r = 0.01 |

For each correlation, p-value and correlation coefficients are shown. When the correlation is significant it is highlighted in bold. Correlations coloured in italics are trends. For each marker, the correlations in the entire cortex (layers I-VI), superficial (I-III) and deep (IV-VI) cortical layers are shown. **Legend**: *Aβ: amyloid beta; DLB: Dementia with Lewy Bodies; LB: Lewy Body; PD: Parkinson's disease; PDD: Parkinson's disease dementia; pSer129-αSyn: phosphorylated Ser129 alpha synuclein; p-NfM/H: phosphorylated neurofilament medium and heavy chain; p-tau: phosphorylated tau.*

**Supplementary Table 6. Correlations of NfL immunoreactivity with pathology load in superficial (layer I-III) and deep cortical layers (layer IV-VI).**

| | **Controls** | **PD + PDD/DLB** | **PD** | **PDD/DLB** |
|---|---|---|---|---|
| ***NfL immunoreactivity (%)*** | | | | |
| *LB count (LB/mm$^2$):* entire cortex | p=0.658, r = 0.05 | **p<0.001, r = 0.36** | p=0.392, r = 0.10 | **p=0.002, r = 0.40** |
| superficial | p=0.851, r = 0.02 | **p<0.001, r = 0.28** | p=0.649, r = 0.05 | **p=0.005, r = 0.33** |
| deep | p=0.144, r = 0.14 | **p=0.005, r = 0.24** | p=0.805, r = 0.02 | **p=0.025, r = 0.27** |
| *pSer129-αSyn load (%):* entire cortex | p=0.666, r = -0.04 | **p<0.001, r = 0.30** | p=0.910, r = 0.01 | **p=0.005, r = 0.36** |
| superficial | p=0.881, r = 0.01 | **p=0.006, r = 0.22** | p=0.997, r = 0.00 | **p=0.021, r = 0.26** |
| deep | p=0.680, r = -0.04 | **p=0.046, r = 0.16** | p=0.217, r = -0.12 | *p=0.074, r = 0.21* |
| *p-tau load (%):* entire cortex | p=0.579, r = -0.06 | **p<0.001, r = 0.42** | **p=0.014, r = 0.31** | **p<0.001, r = 0.42** |
| superficial | p=0.765, r = 0.04 | p=0.112, r = 0.18 | p=0.872, r = 0.02 | p=0.329, r = 0.15 |
| deep | p=0.828, r = 0.03 | p=0.436, r = 0.08 | p=0.802, r = -0.09 | p=0.866, r = 0.03 |
| *Aβ load (%):* entire cortex | p=0.831, r = 0.02 | p=0.499, r = -0.43 | p=0.325, r = -0.09 | p=0.652, r = -0.04 |
| superficial | p=0.461, r = -0.07 | *p=0.051, r = 0.11* | p=0.145, r = -0.10 | *p=0.063, r = 0.15* |
| deep | **p=0.034, r = -0.27** | p=0.924, r = 0.01 | p=0.374, r = 0.11 | p=0.819, r = -0.02 |

For each correlation, p-value and correlation coefficients are shown. When the correlation is significant, it is highlighted in bold, and the strength of the correlation is given by a green shade colour (dark green moderate correlation, light green is a weak correlation). Correlations in italics are trends. For each marker, the correlations in the entire cortex (layers I-VI), superficial (I-III) and deep (IV-VI) cortical layers are shown. **Legend**: *Aβ: amyloid beta; DLB:*

*Dementia with Lewy Bodies; LB: Lewy Body; NfL: neurofilament light chain; PD: Parkinson's disease; PDD: Parkinson's disease dementia; pSer129-αSyn: phosphorylated Ser129 alpha synuclein; p-tau: phosphorylated tau.*

**Supplementary Table 7. Correlations of MRI-derived cortical thickness with neurofilaments immunoreactivity and pathology load.**

| | Controls | PD + PDD/DLB | PD | PDD/DLB |
|---|---|---|---|---|
| ***Cortical thickness (Z score)*** | | | | |
| *p-NfM/H immunoreactivity (%)* | p=0.686, r = 0.04 | **p=0.020, r = -0.23** | **p=0.047, r = -0.32** | p=0.106, r = -0.21 |
| *NfL immunoreactivity (%)* | p=0.680, r = 0.33 | **p=0.012, r = -0.22** | **p=0.025, r = -0.27** | p=0.220, r = -0.15 |
| *LB count (LB/mm$^2$)* | - | p=0.910, r = -0.01 | p=0.506, r = -0.08 | p=0.615, r = -0.05 |
| *pSer129-αSyn load (%)* | - | p=0.784, r = 0.02 | p=0.781, r = -0.04 | p=0.405, r = 0.09 |
| *p-tau load (%)* | p=0.400, r = 0.06 | p=0.106, r = -0.13 | *p=0.091, r = -0.20* | p=0.440, r = -0.08 |
| *Aβ load (%)* | p=0.488, r = 0.68 | p=0.876, r = -0.01 | p=0.632, r = -0.06 | p=0.367, r = 0.12 |

For each correlation, p-value and correlation coefficients are shown. When the correlation is significant it is highlighted in bold, and the strength of the correlation is given by a green shade colour (dark green moderate correlation, light green is a weak correlation). Correlations in italics are trends. **Legend:** *Aβ: amyloid beta; DLB: Dementia with Lewy Bodies; LB: Lewy Body; PD: Parkinson's disease; PDD: Parkinson's disease dementia; pSer129-αSyn: phosphorylated Ser129 alpha synuclein; p-NfM/H: phosphorylated neurofilament medium and heavy chain; p-tau: phosphorylated tau.*

**Supplementary Table 8. Correlations of DTI-derived cortical MD with neurofilaments immunoreactivity and pathology load.**

| | **Controls** | **PD + PDD/DLB** | **PD** | **PDD/DLB** |
|---|---|---|---|---|
| ***FWC-MD ($10^{-3}$ $mm^2$/s)*** | | | | |
| *p-NfM/H immunoreactivity (%)* | p=0.430, r = -0.08 | p=0.182, r = 0.14 | p=0.586, r = 0.10 | **p=0.003, r = 0.28** |
| *NfL immunoreactivity (%)* | p=0.738, r = 0.03 | **p=0.024, r = 0.21** | p=0.184, r = -0.19 | **p=0.003, r = 0.41** |
| *LB count (LB/$mm^2$)* | - | **p=0.002, r = 0.23** | p=0.533, r = -0.07 | **p=0.002, r = 0.32** |
| *pSer129-αSyn load (%)* | - | **p=0.003, r = 0.23** | p=0.874, r = 0.02 | **p=0.002, r = 0.32** |
| *p-tau load (%)* | **p=0.003, r = 0.23** | **p<0.001, r = 0.28** | p=0.393, r = 0.10 | **p<0.001, r = 0.36** |
| *Aβ load (%)* | p=0.779, r = -0.01 | p=0.850, r = 0.02 | p=0.366, r = -0.11 | p=0.339, r = 0.00 |

For each correlation, p-value and correlation coefficients are shown. When the correlation is significant it is highlighted in bold, and the strength of the correlation is given by a green shade colour (dark green moderate correlation, light green is a weak correlation). **Legend**: *Aβ: amyloid beta; DLB: Dementia with Lewy Bodies; DTI: diffusion tensor imaging; FWC-MD: free-water mean diffusivity; LB: Lewy Body; PD: Parkinson's disease; PDD: Parkinson's disease dementia; pSer129-αSyn: phosphorylated Ser129 alpha synuclein; p-NfM/H: phosphorylated neurofilament medium and heavy chain; p-tau: phosphorylated tau.*

### References supplementary tables


1 Braak H, Alafuzoff I, Arzberger T, Kretzschmar H, Del Tredici K (2006) Staging of Alzheimer disease-associated neurofibrillary pathology using paraffin sections and immunocytochemistry. Acta neuropathologica 112: 389-404

2 Braak H, Del Tredici K, Rub U, de Vos RA, Jansen Steur EN, Braak E (2003) Staging of brain pathology related to sporadic Parkinson's disease. Neurobiol Aging 24: 197-211 Doi 10.1016/s0197-4580(02)00065-9

3 Daducci A, Gerhard S, Griffa A, Lemkaddem A, Cammoun L, Gigandet X, Meuli R, Hagmann P, Thiran JP (2012) The connectome mapper: an open-source processing pipeline to map connectomes with MRI. PLoS One 7: e48121 Doi 10.1371/journal.pone.0048121

4 Hagmann P, Cammoun L, Gigandet X, Meuli R, Honey CJ, Wedeen VJ, Sporns O (2008) Mapping the structural core of human cerebral cortex. PLoS Biol 6: e159 Doi 10.1371/journal.pbio.0060159

5 Montine TJ, Phelps CH, Beach TG, Bigio EH, Cairns NJ, Dickson DW, Duyckaerts C, Frosch MP, Masliah E, Mirra SS (2012) National Institute on Aging–Alzheimer's Association guidelines for the neuropathologic assessment of Alzheimer's disease: a practical approach. Acta neuropathologica 123: 1-11

6 Thal DR, Rüb U, Orantes M, Braak H (2002) Phases of Aβ-deposition in the human brain and its relevance for the development of AD. Neurology 58: 1791-1800
